# Predicting enhancer-gene links from single-cell multi-omics data by integrating prior Hi-C information

**DOI:** 10.1101/2025.10.09.681330

**Authors:** Xuan Liang, Yuanyuan Miao, Dongmei Han, Yurun Li, Wenwen Zhang, Zhen Wang

**Affiliations:** Shanghai Institute of Nutrition and Health, University of Chinese Academy of Sciences, Chinese Academy of Sciences, Shanghai 200031, China; Key Laboratory of Systems Health Science of Zhejiang Province, School of Life Science, Hangzhou Institute for Advanced Study, University of Chinese Academy of Sciences, Hangzhou 310024, China

**Keywords:** single-cell multi-omics, bulk average Hi-C, weighted graphical lasso, enhancer-gene links

## Abstract

Enhancers play an important role in transcriptional regulation by modulating gene expression from distal genomic locations. Although single-cell ATAC and RNA sequencing (scATAC/RNA-seq) data have been leveraged to infer enhancer-gene links, establishing regulatory links between enhancers and their target genes remains a challenge due to the absence of chromatin conformation information. Here, we present SCEG-HiC, a machine learning method based on weighted graphical lasso, which decodes enhancer-gene links from single-cell multi-omics data by integrating bulk average Hi-C as prior knowledge. Comprehensive evaluation across ten single-cell multi-omics datasets from both humans and mice demonstrates that SCEG-HiC outperforms existing single-cell models, regardless of using paired scATAC/RNA-seq or scATAC-seq data alone. Application of SCEG-HiC to COVID-19 datasets illustrates its capacity to more reliably reconstruct gene regulatory networks underlying disease severity, and elucidate functional associations between non-coding variants and their putative target genes. SCEG-HiC is freely available as an open-source and user-friendly R package, facilitating broad applications in regulatory genomics research.

## Introduction

In eukaryotes, transcriptional regulation is an essential biological process that enables cells or organisms to respond to various intra- and extra-cellular signals, maintain differentiated cell types, and coordinate cellular activities^1^. Promoters and enhancers, two major types of *cis*-regulatory elements (CREs), are short non-coding sequences bound by transcription factors (TFs) to govern transcriptional regulation of adjacent genes^2, 3^. While promoters initiate transcription at or near the transcription start site (TSS), enhancers can augment transcription from more distal genomic locations. Genome-wide association studies (GWAS) have shown that > 90% of single nucleotide polymorphisms (SNPs) associated with complex diseases are in non-coding regions of the genome far from promoters, implicating a potential regulatory role mediated by enhancers^4^. However, linking enhancers and their target genes (or equivalently, promoters) remains a significant challenge. First, enhancers are position-independent; they can be located upstream or downstream of their target promoter. Multiple studies have shown that enhancers can regulate genes located hundreds of thousands to millions of base pairs away, and they often do not regulate the closest gene^5–7^. Second, enhancers regulate the target gene expression in a cell- and tissue-specific manner, meaning that the same enhancer may target different genes under different biological contexts^3^. Third, enhancer-gene relationships are many-to-many: one enhancer can regulate multiple genes, and one gene may be controlled by multiple enhancers^8^.

Several experimental approaches have been developed to identify enhancer-gene links. Chromatin conformation capture (3C)^9^ and its derivative genomic assays, such as high-throughput chromatin conformation capture (Hi-C)^10^, detect physical interactions between genomic regions. Other approaches, such as those based on expression quantitative trait loci (eQTL)^11^ and CRISPR-Cas9 perturbation assays^12^, infer regulatory links by associating genetic variants in enhancer regions with gene expression change. However, these assays are often technically challenging and costly, and have only been conducted at high resolution in a limited range of cell types.

As a supplement to experimental methods, various computational methods have been proposed. The simplest approach assigns an enhancer to its closest gene, but it overlooks the fact that enhancers can regulate gene expression over long genomic distances. As CRE activities can be measured via histone modification or chromatin accessibility profiling (e.g., H3K27ac ChIP-seq, ATAC-seq), several studies have modeled multi-omics signals from diverse tissues and cell types to infer enhancer-gene links^13–15^. Correlation-based methods calculate the statistical correlation of activity levels between enhancers and gene promoters^16, 17^. Supervised learning-based methods, such as IM-PET^18^, TargetFinder^19^, and McEnhancer^20^, build linear or non-linear classifier based on a set of verified enhancer-gene links. Score-based methods integrate multiple types of information to generate quantitative scores for assigning target genes to enhancers^21, 22^. For instance, the activity-by-contact (ABC) model combines enhancer activity with chromatin contact data (Hi-C) to compute quantitative scores^23^. Notably, the ABC model outperformed several correlation- and supervised learning-based methods and showed comparable performance when using either cell-type specific Hi-C or average Hi-C data across multiple cell types. Yet, bulk sequencing technologies measure the average signals across biological tissues, underestimating cellular heterogeneity and lacking the ability to capture cell- and state-specific information.

The advent of single-cell technologies, such as single-cell RNA sequencing (scRNA-seq)^24^ and single-cell ATAC sequencing (scATAC-seq)^25^, has made it possible to uncover cellular heterogeneity at single-cell resolution. Notably, techniques such as SHARE-seq^26^ and 10x Genomics Multiome ATAC + RNA^27^ enable simultaneous measurement of gene expression and chromatin accessibility within the same cell. In principle, computational strategies developed for bulk sequencing data can be adopted for single-cell sequencing data by replacing bulk samples with individual cells^28^. For example, Signac^29^ and FigR^30^ use Pearson and Spearman correlation coefficients between gene expression and chromatin accessibility to identify enhancer-gene links, respectively. SCNEIC+^31^ and DIRECT-NET^32^ build machine learning models to predict gene expression using a combination of nearby peaks and extract enhancer-gene links based on feature importance scores. Cicero, a method designed to identify co-accessible peaks solely using scATAC-seq data, regularizes the correlation matrix via the graphic lasso (glasso) algorithm to eliminate indirect links^33^. Cicero also penalizes partial correlations in a distant-dependent manner, where peaks closer in genomic distance have a lower penalty term. Nonetheless, previous studies largely focused on gene expression and chromatin accessibility, ignoring chromatin confirmation in single cells. Although single-cell Hi-C (scHi-C)^34, 35^ techniques have been developed, such data are scarce due to their experimental complexity. To overcome this, we hypothesized that bulk Hi-C data averaged over multiple cell types can also serve as a proxy in single-cell contexts, as validated by the ABC model^23^.

Here, we propose SCEG-HiC, a computational method that predicts **E**nhancer-**G**ene links from **S**ingle-**C**ell multi-omics data by integrating prior Hi-C information. SCEG-HiC employs the weighted graphical lasso (wglasso) model^36^ to incorporate average bulk Hi-C data, effectively regularizing the correlation matrix with the prior Hi-C contact matrix as a penalty term. SCEG-HiC can use either scATAC-seq alone or paired scATAC/RNA-seq data to predict regulatory relationships. Through extensive benchmarking using independent functional genomics data including cell-type-specific Hi-C and eQTL, we demonstrate that SCEG-HiC consistently outperforms existing enhancer-gene linking models across diverse human and mouse datasets. Moreover, we illustrate the biological relevance of SCEG-HiC by constructing cell-type-specific gene regulatory networks (GRNs) and prioritizing target genes associated with non-coding GWAS variants. SCEG-HiC is implemented as an R package and is freely available at: https://github.com/wuwei77lx/SCEGHiC.

## Results

### Overview of SCEG-HiC framework

SCEG-HiC is a computational framework designed to predict enhancer-gene links using single-cell multi-omics data (paired scATAC/RNA-seq or scATAC-seq alone), combined with three-dimensional chromatin conformation information derived from bulk average Hi-C data. Gene promoters are defined as ± 1 kb regions around TSS, while candidate enhancers are identified as accessible chromatin regions outside promoters but within a user-specified genomic window (default: ± 250 kb). The SCEG-HiC framework comprises three major steps (Fig. 1 and Methods): (i) preprocessing CRE accessibility and gene expression matrices; (ii) constructing and normalizing a bulk average Hi-C contact matrix; (iii) inferring enhancer-gene links using the wglasso model.

**Fig. 1.**
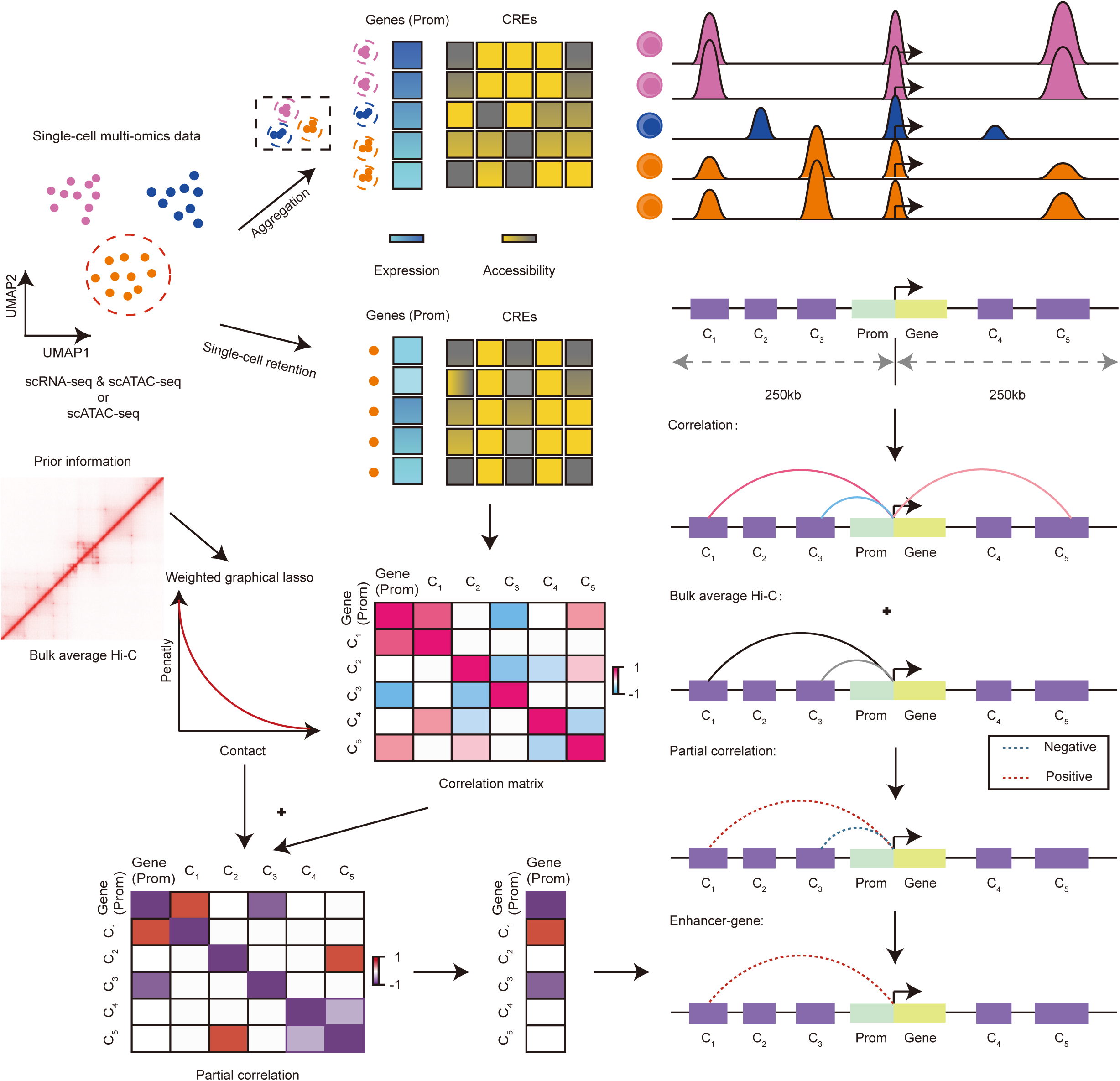
The workflow of SCEG-HiC. SCEG-HiC integrates single-cell multi-omics data and bulk average Hi-C data to infer enhancer-gene links. The left panel illustrates the detailed workflow of SCEG-HiC. Taking paired scATAC/RNA-seq or scATAC-seq data alone as input, SCEG-HiC first computes enhancer-gene correlation matrices based on gene expression (or promoter accessibility) and distal CRE accessibility across aggregated cells or single cells from a specific cell type. Second, it incorporates chromatin interaction priors from bulk average Hi-C data to construct a normalized contact matrix between genes and CREs. Third, SCEG-HiC applies a wglasso model to estimate partial correlation coefficients, preserving enhancer-gene links with positive coefficients. The right panel shows a schematic genomic region illustrating how SCEG-HiC integrates Hi-C priors with single-cell omics correlations to calculate partial correlation coefficients for enhancer-gene links. For each gene (green), the ± 1 kb region around the TSS is defined as the promoter (cyan), while CREs within ± 250 kb regions (excluding promoters) are defined as candidate enhancers (purple, e.g., C2, C3, and C4). Prom: promoter.

The preprocessing step mainly involves two considerations: the sparse nature of single-cell sequencing data (especially scATAC-seq), and spurious peak-gene associations confounded by cell type heterogeneity^37, 38^. To systematically evaluate these issues, we compared six preprocessing strategies (Supplementary Fig. 1, Supplementary Table 1 and Methods), considering either aggregation or single-cell retention approach, along with the impact of binarizing scATAC-seq data. The aggregation approach, applied across cell types, aggregates signals from small clusters of similar cells to alleviate data sparsity. The single-cell retention approach, tailored to specific cell types, preserves individual cell signals. Our results showed that the single-cell retention strategy with normalized scATAC-seq data achieved the highest prediction accuracy, albeit identifying fewer enhancer-gene links. In contrast, the aggregation approach with binarized scATAC-seq data resulted in slightly lower accuracy but captured a broader range of enhancer-gene links. To balance accuracy and coverage, we implemented both preprocessing strategies in SCEG-HiC, with aggregation designated as the default.

Distinct from previous methods, SCEG-HiC integrates prior knowledge of chromatin conformation to improve prediction accuracy. Although chromatin interactions can be cell-type specific, previous studies have shown that many chromatin interactions are highly similar across different cell types^39^. Notably, an average Hi-C map generated from diverse cell types can capture potential chromatin interactions in new cell types, as validated by the ABC model^23^ (Methods). Due to the low contact frequency in bulk average Hi-C data, we performed normalization on the contact matrix. Among four evaluated normalization methods, the rank score method significantly outperformed the others in terms of prediction accuracy (Supplementary Fig. 2, Supplementary Table 1 and Methods).

Finally, SCEG-HiC employs the wglasso model to infer enhancer-gene links, effectively addressing the high-dimensionality of single-cell data while integrating prior information^36^. Specifically, SCEG-HiC integrates the normalized Hi-C contact matrix with the covariance matrix of gene expression (or promoter accessibility) and candidate enhancer accessibility. The penalty parameter is optimized using Bayesian information criterion (BIC) cross-validation to estimate the inverse covariance matrix^40^. Partial correlation coefficients extracted from this matrix quantify the strength of enhancer-gene links, with positive values indicating biologically meaningful links.

### Benchmarking SCEG-HiC with human paired single-cell multi-omics data

We analyzed five paired scATAC/RNA-seq datasets from different human tissues generated by 10x Genomics, including peripheral blood mononuclear cells (PBMCs), skin stromal cells^41^, fetal retinal tissue^42^, brain gray matter^43^, and developing cerebral cortex^44^ (Fig. 2a and Supplementary Table 2). For each dataset, we applied Seurat^45^ and Signac^29^ to perform quality control (QC), cell clustering, cell type annotation, and uniform manifold approximation and projection (UMAP) visualization based on weighted nearest neighbor (WNN) integration of gene expression and chromatin accessibility data (Fig. 2a, Supplementary Fig. 3 and Methods). We applied SCEG-HiC to each dataset to construct enhancer-gene links for marker genes of each cell type. To facilitate consistent comparison with other methods, here we utilized aggregation approach with binarized scATAC-seq data. The bulk average Hi-C data for humans were obtained from the ABC model and preprocessed across ten cell lines (GM12878, NHEK, HMEC, RPE1, THP1, IMR90, HUVEC, HCT116, K562, and KBM7)^46^.

**Fig. 2.**
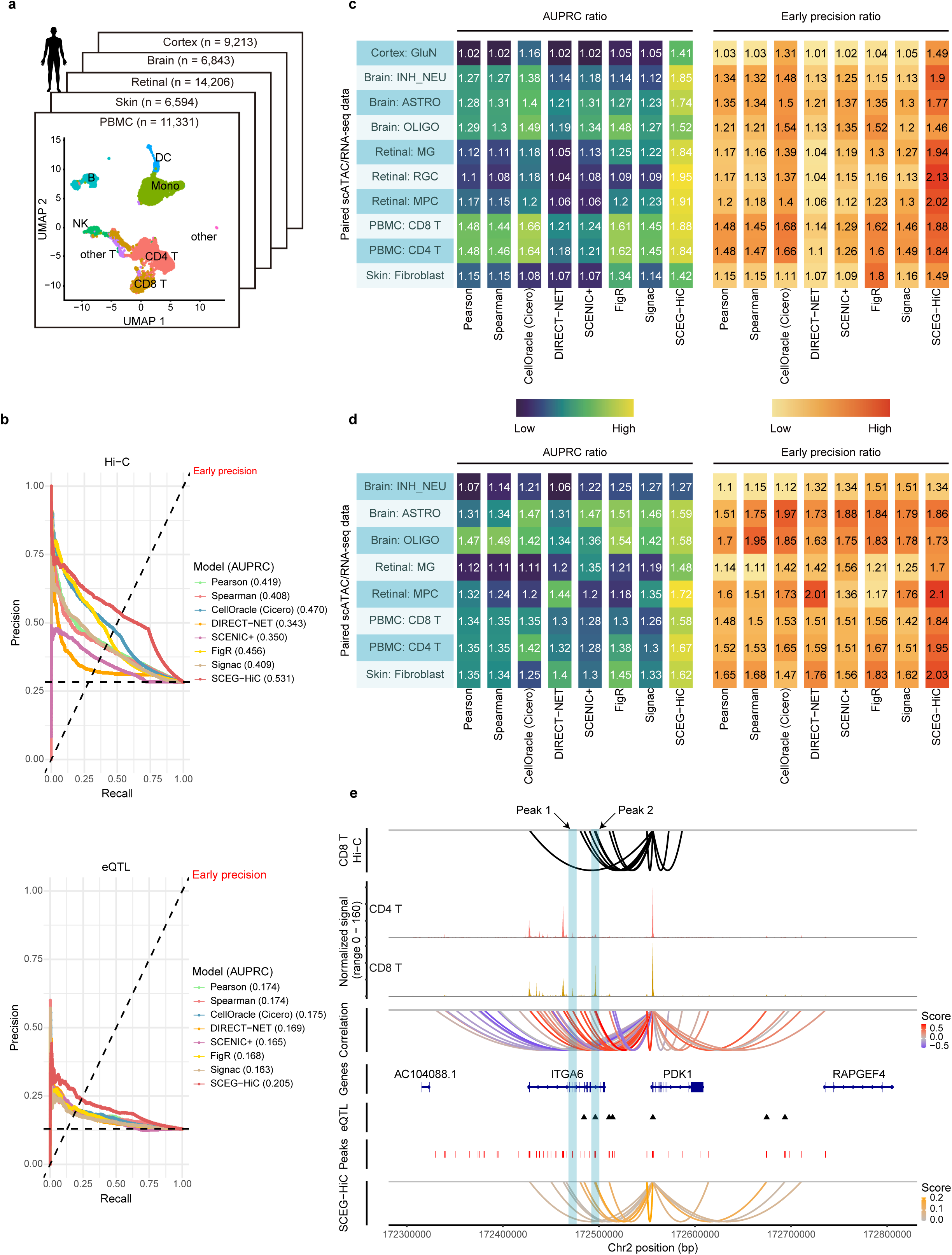
SCEG-HiC can accurately predict enhancer-gene links using human paired single-cell multi-omics data. **a**, Summary of human paired scATAC/RNA-seq datasets used for benchmarking (*n* = cell number), along with UMAP visualizations of a representative PBMC dataset. Cell types: CD4 T, CD4+ T cell; CD8 T, CD8+ T cell; Mono, monocyte; DC, dendritic cell; NK, natural killer cell; other T: other T cell. Datasets: skin, skin stromal cells; retinal, fetal retinal tissue; brain, brain gray matter; cortex, developing cerebral cortex. **b**, Precision-recall curves evaluating enhancer-gene link predictions performance for CD8+ T cells, validated using CD8+ T cell-specific Hi-C (top) and whole blood eQTL data (bottom). Models compared include SCEG-HiC, CellOracle (Cicero-based), DIRECT-NET, SCENIC+, FigR, Signac, and correlation methods (Pearson, Spearman). The intersection of the dashed diagonal line with the precision-recall curve indicates EP, while the horizontal dashed line represents the random prediction baseline. AUPRC, area under the precision-recall curve. **c-d**, Performance of enhancer-gene link predictions across different human cell types, measured by AUPRC ratio (left) and EPR (right). Validation was performed using cell type-specific Hi-C **(c)** and eQTL data **(d)**, respectively. **e**, Representative example of predicted enhancer-gene links for *PDK1*. Integration of CD8+ T cell Hi-C interactions, scATAC-seq signals, aggregated ATAC-RNA correlations, eQTL SNPs, and enhancer-gene links predicted by SCEG-HiC around *PDK1*. Peak 1 represents an enhancer not predicted by SCEG-HiC, exhibiting strong correlations but lacking Hi-C and eQTL support. Peak 2 corresponds to an enhancer predicted by SCEG-HiC, supported by both Hi-C and eQTL evidence.

To evaluate the accuracy of predicting enhancer-gene links in the human paired single-cell multi-omics data, we compared SCEG-HiC against two classical correlation-based methods (Pearson and Spearman), and five dedicated computational tools: CellOracle^47^, DIRECT-NET^32^, SCENIC+^31^, FigR^30^, and Signac^29^ (Methods). We collected experimentally supported interactions from cell type-specific Hi-C and eQTL data as the benchmarks. Given the highly imbalanced fraction of verified enhancer-gene interactions, we used the area under the precision-recall curve (AUPRC) and early precision (EP) as evaluation metrics, which are inherently threshold-independent^48, 49^ (Methods). Using CD8+ T cells from the PBMC dataset, we first validated the performance of SCEG-HiC against CD8+ T cell Hi-C data from ENCODE^50^ and whole blood eQTL data from GTEx^51^. In Hi-C-based validation, SCEG-HiC achieved an AUPRC of 0.531 and an EP of 0.531; in eQTL-based validation, it reached an AUPRC of 0.205 and an EP of 0.238, substantially outperforming all other methods (Fig. 2b).

To further illustrate the broad applicability of SCEG-HiC, we evaluated its performance across nine additional representative cell types from the five multi-omics datasets. These included cell types covered and uncovered by the bulk average Hi-C data, allowing a comprehensive assessment of model robustness across diverse cellular contexts. To ensure comparability across datasets and cell types, we calculated AUPRC ratio and EP ratio (EPR) relative to the fraction of verified interactions (i.e., random baseline) for each cell type within each dataset. Using Hi-C-validated enhancer-gene links from all ten cell types, SCEG-HiC consistently yielded significantly higher AUPRC ratio and EPR than other methods across most cases (Fig. 2c). A similar trend was observed when using eQTL-derived links from eight cell types as benchmarks: SCEG-HiC again outperformed other methods in terms of both AUPRC ratio and EPR across most cases (Fig. 2d). Supplementary Table 3 includes *P*-values and 95% confidence intervals from paired *t*-tests or Wilcoxon tests comparing SCEG-HiC to other models. These evaluation results collectively confirmed that SCEG-HiC achieves superior performance in predicting enhancer-gene links from human paired single-cell multi-omics data.

To further confirm that the improvement of SCEG-HiC stemmed from the incorporation of bulk average Hi-C data, we performed an ablation analysis comparing SCEG-HiC (with Hi-C data) and SCEG-Base (without Hi-C data) under the same model framework. Across both cell type-specific Hi-C and eQTL validation datasets, SCEG-HiC consistently outperformed SCEG-Base in terms of AUPRC ratio (Hi-C: *P* = 3.809 × 10^−6^, eQTL: *P* = 1.090 × 10^−6^ via two-sided paired *t*-test; Supplementary Fig. 4a) and EPR (Hi-C: *P* = 8.381 × 10^−6^, eQTL: *P* = 1.043 × 10^−4^ via two-sided paired *t*-test; Supplementary Fig. 4b). These results confirmed that the performance gains of SCEG-HiC are indeed primarily attributed to the integration of Hi-C data.

As a case study, we examined enhancer-gene links involving *PDK1*, a key regulator of T cell development and activation, using the PBMC dataset (Fig. 2e)^52, 53^. SCEG-HiC accurately recovered validated links using CD8+ T cell-specific Hi-C data from ENCODE and whole blood eQTL data from GTEx. Importantly, we observed that integrating bulk average Hi-C information substantially reduces false positives compared to models relying solely on correlations between gene expression and chromatin accessibility.

### Benchmarking SCEG-HiC with human scATAC-seq data alone

To assess the accuracy of predicting enhancer-gene links using human scATAC-seq data alone, we compared SCEG-HiC with two statistical methods, Pearson correlation and chi-squared test with false discovery rate correction (Chi2+FDR), as well as three computational tools: Cicero^33^, DIRECT-NET^32^, and scEchIA^54^ (Methods). DIRECT-NET is suitable for both parallel scATAC/RNA-seq data and scATAC-seq data alone. scEchIA shares similarities with our approach, using average chromatin interactions from other cell types to predict links between distal sites (25 kb) in a specific cell type. Using the same five paired single-cell multi-omics datasets, we restricted our analysis to the scATAC-seq data to predict enhancer-gene links. Data preprocessing and integration of prior bulk average Hi-C data in SCEG-HiC were consistent with the approach used for paired multi-omics data.

Similarly, we validated the inferred enhancer-gene links against both cell type-specific Hi-C and eQTL data, using AUPRC ratio and EPR as evaluation metrics (Methods). In Hi-C-based validation across ten cell types, SCEG-HiC significantly outperformed all other methods in both AUPRC ratio and EPR (Fig. 3a and Supplementary Table 3). Notably, scEchIA demonstrated the second-best performance, closely following SCEG-HiC. For instance, in the fetal retinal tissue dataset, the AUPRC ratio and EPR achieved by scEchIA were only slightly lower than those of SCEG-HiC, yet markedly higher than those of the other models. This observation further supports the notion that incorporating chromatin interaction information can substantially enhance model prediction accuracy. In eQTL-based validation across eight cell types, SCEG-HiC again showed significantly higher AUPRC ratio and EPR across most cell types (Fig. 3b and Supplementary Table 3). However, in contrast to the Hi-C validation results, scEchIA did not consistently rank second in accuracy, possibly due to its lack of Hi-C data from a broader set of cell types. In summary, these results demonstrate that SCEG-HiC maintains robust predictive performance when operating solely on human scATAC-seq data.

**Fig. 3.**
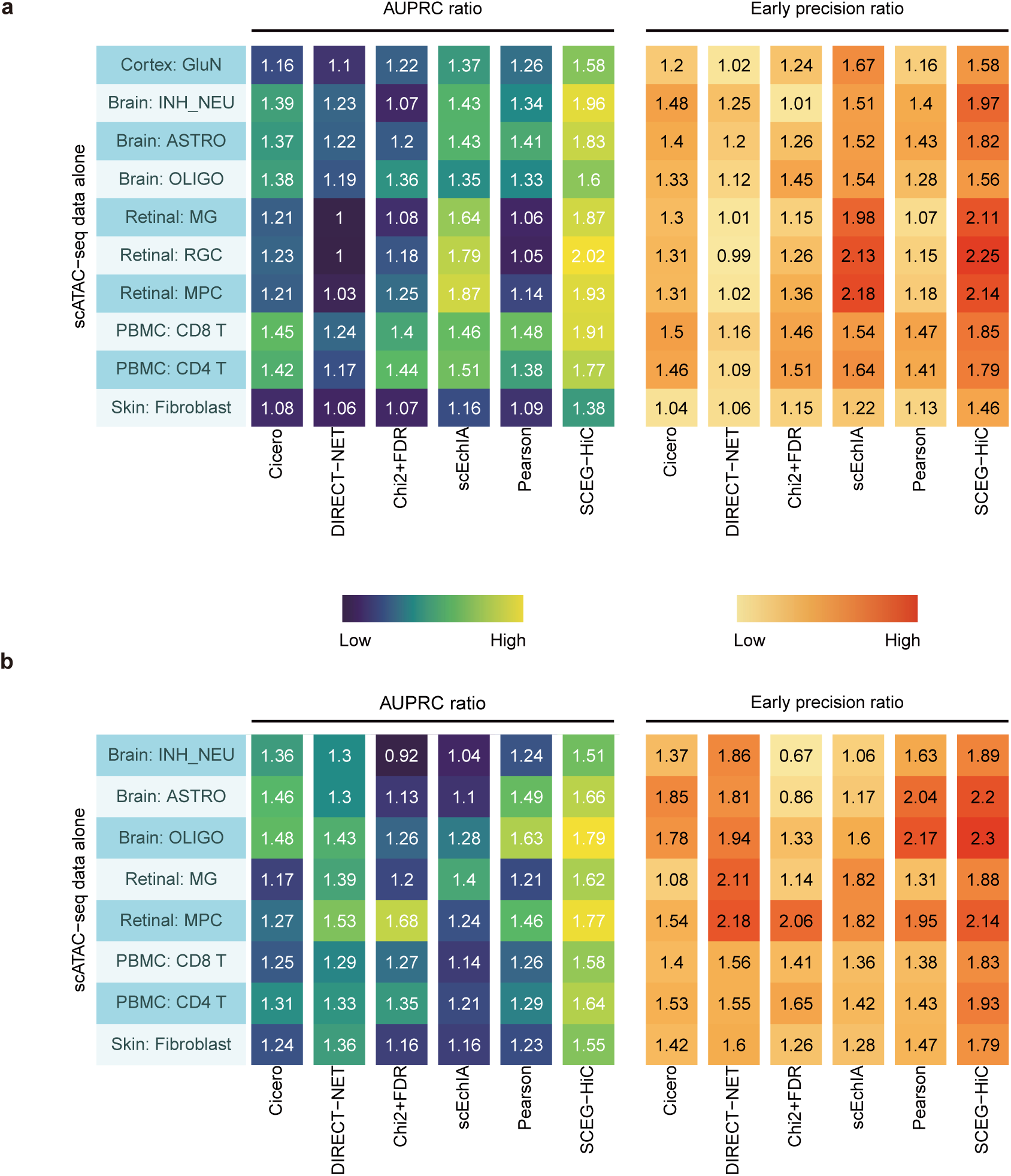
SCEG-HiC can accurately predict enhancer-gene links using human scATAC-seq data alone. **a-b**, Performance of enhancer-gene link predictions across different human cell types, measured by AUPRC ratio (left) and EPR (right). Validation was performed using cell type-specific Hi-C **(a)** and eQTL data **(b)**. Five competing methods were considered for comparison, including Cicero, DIRECT-NET, Chi2+FDR, scEchIA, and Pearson correlation.

### Benchmarking SCEG-HiC with mouse paired single-cell multi-omics data

We analyzed five paired single-cell multi-omics datasets from different mouse tissues generated by different platforms, including mouse embryonic brain cells at day 18 (10x Genomics), mouse skin (SHARE-seq)^26^, adult mouse cerebral cortex (SHARE-seq)^55^, thymic epithelial cells (10x Genomics)^56^, and mouse liver (10x Genomics)^57^ (Fig. 4a, and Supplementary Table 2). Each dataset was processed using Seurat and Signac for QC, dimensionality reduction, UMAP visualization, clustering and cell annotation (Fig. 4a, Supplementary Fig. 5 and Methods). Given the absence of pre-existing average Hi-C data for mice, we collected seven Hi-C datasets from the ENCODE project, including two embryonic stem cell lines (mESC1, mESC2), CH12LX, CH12F3, fiber, epithelium, and B cells^50^ (Supplementary Table 4). Following the ABC model’s approach for constructing human average Hi-C data, we scaled each Hi-C dataset using dataset-specific power-law distributions and computed their average to generate the mouse average Hi-C data (Fig. 4b and Methods). Subsequently, we applied SCEG-HiC to each mouse single-cell dataset with aggregation and scATAC-seq binarization approach, integrating the constructed mouse average Hi-C data to predict enhancer-gene links.

**Fig. 4.**
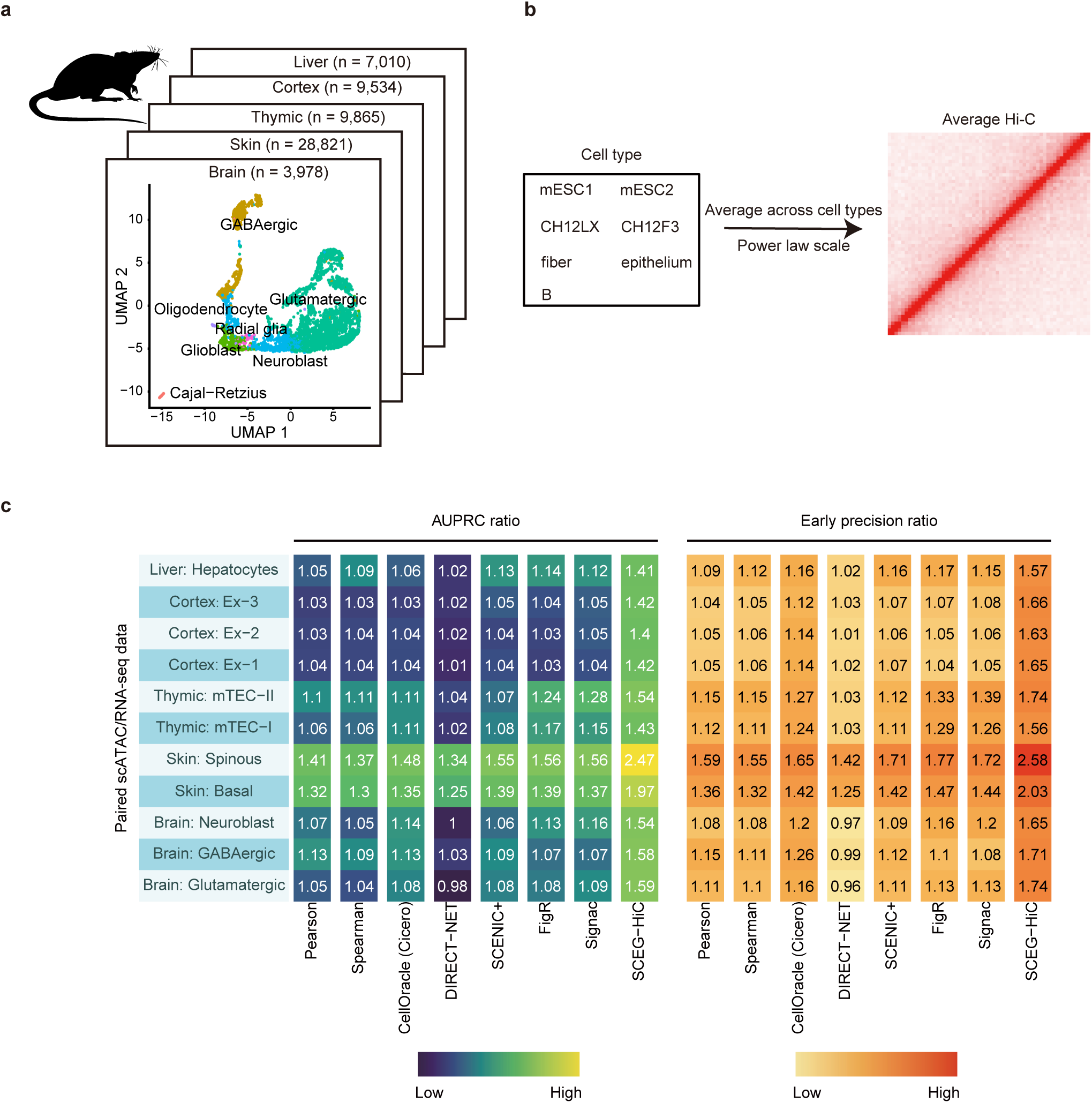
SCEG-HiC can accurately predict enhancer-gene links using mouse paired single-cell multi-omics data. **a**, Summary of mouse paired scATAC/RNA-seq datasets used for benchmarking (*n* = cell number), along with UMAP visualizations of a representative mouse embryonic brain cells at day 18 dataset. Datasets: brain, mouse embryonic brain cells at day 18; skin, mouse skin; thymic, thymic epithelial cells; cortex, adult mouse cerebral cortex; liver, mouse liver. **b**, A schematic illustration for generating mouse bulk averaged Hi-C data from seven datasets. **c**, Performance of enhancer-gene link predictions across different mouse cell types, assessed by AUPRC ratio (left) and EPR (right) using cell type-specific Hi-C data. Seven competing methods were considered for comparison, including Pearson correlation, Spearman correlation, CellOracle (Cicero-based), DIRECT-NET, SCENIC+, FigR, and Signac.

We benchmarked SCEG-HiC against seven methods or tools for paired scATAC/RNA-seq data, as mentioned before. Due to the lack of eQTL data for mice, we only used cell type-specific Hi-C data for benchmarks, with AUPRC ratio and EPR as evaluation metrics. Across the five datasets, we selected eleven representative cell types for evaluation, including those covered and uncovered by the mouse bulk average Hi-C data. We observed that SCEG-HiC consistently achieved significantly higher AUPRC ratio and EPR compared to all other methods (Fig. 4c and Supplementary Table 3). Furthermore, the performance improvements observed in mouse datasets were even more pronounced than those in human datasets, highlighting the robustness and cross-species applicability of SCEG-HiC. In conclusion, we validated SCEG-HiC as a superior method in predicting enhancer-gene links using mouse paired single-cell multi-omics data.

### Comparison of SCEG-HiC and ABC

Although the ABC model was originally designed for bulk sequencing data, it has also been adopted in scATAC-seq analysis by aggregating cells into pseudobulk profiles. Given that both SCEG-HiC and ABC utilize bulk average Hi-C data, we systematically compared the two models to evaluate the improvements attained by SCEG-HiC^23^. First, we assessed the prediction accuracy using both cell type-specific Hi-C and eQTL data. In Hi-C-based validation, the ABC model showed a significantly higher AUPRC ratio in both human and mouse datasets (human: *P* = 0.007, mouse: *P* = 9.766 × 10^−4^ via two-sided paired *t*-test). In contrast, SCEG-HiC consistently outperformed the ABC model in terms of EPR (human: *P* = 2.138 × 10^-s^, mouse: *P* = 1.246 × 10^−4^ via two-sided paired *t*-test; Fig. 5a-d). In eQTL-based validation, no significant differences were observed between the two models in terms of AUPRC ratio (*P* = 0.069 via two-sided paired *t*-test), though the ABC model demonstrated superior performance in the EPR metric (*P* = 0.040 via two-sided paired *t*-test).

**Fig. 5.**
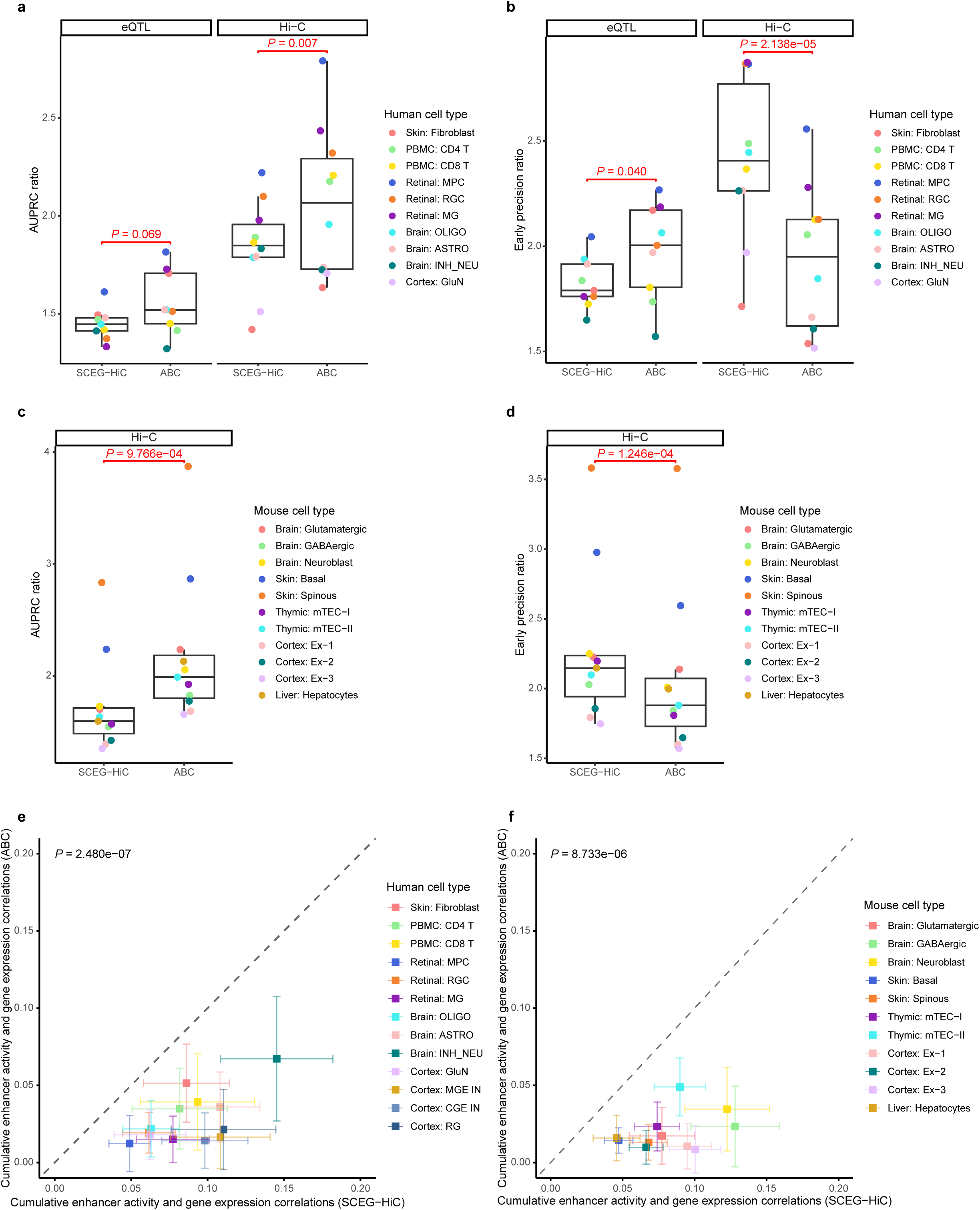
SCEG-HiC demonstrates higher expression consistency compared to ABC. **a-d**, Boxplots comparing performance between SCEG-HiC and ABC across various human **(a, b)** and mouse **(c, d)** cell types, measured by AUPRC ratio **(a, c)** and EPR **(b, d)**. Validation was performed using cell type-specific Hi-C data for both species, and eQTL data for humans only. Each dot represents a distinct cell type. *P*-values were calculated using two-sided paired *t*-tests. **e-f**, Cumulative enhancer activity and gene expression correlations compared between SCEG-HiC and ABC across human **(e)** and mouse **(f)** cell types. The x- and y-axes represent Spearman correlations from SCEG-HiC and ABC predictions, respectively. Each dot corresponds to the mean correlation across genes for a specific cell type, with error bars representing ± 0.5 × standard deviation. The dashed diagonal line indicates equal correlations, and *P*-values were calculated between the two models using two-sided paired *t*-tests.

Next, considering that gene expression is typically regulated by the combined activity of multiple enhancers, we evaluated the functional relevance of predicted enhancer-gene links from both models. Specifically, we summed the ATAC-seq signals across all enhancers associated with each gene at the single-cell level, then calculated the Spearman correlations between the cumulative enhancer activity and corresponding gene expression (Methods). This metric assesses the functional plausibility of predicted enhancer-gene links, under the assumption that accurate regulatory relationships should manifest as a strong positive correlation between cumulative enhancer activity and target gene expression. Across diverse human and mouse cell types, SCEG-HiC consistently outperformed the ABC model, showing significantly stronger correlations between predicted cumulative enhancer activity and gene expression (human: *P* = 2.480 × 10^−7^, mouse: *P* = 8.733 × 10^−6^ via two-sided paired *t*-test; Fig. 5e, f and Supplementary Fig. 6). This expression concordance analysis highlights a critical distinction between single-cell methods that account for cell-to-cell variability and the ABC model, which does not, thereby complementing validations based on Hi-C or eQTL data.

In summary, ABC and SCEG-HiC exhibit relative advantages in Hi-C and eQTL-based benchmarks, with performance dependent on specific metrics. Notably, SCEG-HiC demonstrates significantly stronger correlations between predicted cumulative enhancer activity and gene expression, a key advantage that facilitates regulatory interpretability of predicted enhancer-gene links.

### Analysis of COVID-19 unpaired single-cell multi-omics data

We applied SCEG-HiC to unpaired PBMC scRNA-seq and scATAC-seq datasets from patients infected with SARS-CoV-2, focusing on enhancer-mediated regulations underlying severe and mild COVID-19 symptoms^58^. The patients were stratified into mild and severe categories based on the World Health Organization (WHO) severity scores (severe: 5-7, mild: 2-4). We utilized the scRNA-seq data preprocessed and clustered by Wilk et al., selecting 18 samples that met our study requirements. This dataset comprised a total of 61,486 cells spanning 14 distinct cell types (Supplementary Fig. 7a-c and Methods). For scATAC-seq analysis, we selected 17 COVID-19 samples (9 mild and 8 severe), and processed the data using Signac. After QC, we retained 52,817 high-quality cells, performed peak calling on each sample using MACS2, and generated a unified peak set comprising 195,424 peaks^59^. Following data integration, clustering, and annotation, 10 cell types were identified (Supplementary Fig. 7d-f and Methods). Differential expression analysis per cell type between mild and severe groups revealed that CD14+ monocytes exhibited the highest number of differentially expressed genes (DEGs), highlighting this cell type’s pronounced response to SARS-CoV-2 infection (Supplementary Fig. 8a and Methods). In total, 871 genes showed significant differential expression in CD14+ monocytes (|log2-fold change (FC)| > 0.25, adjusted *P* < 0.05; Fig. 6a). We then applied SCEG-HiC to the scATAC-seq data of CD14+ monocytes using single-cell retention approach and benchmarked its performance against Cicero, a classic method that predicts enhancer–gene links using scATAC-seq data alone.

**Fig. 6.**
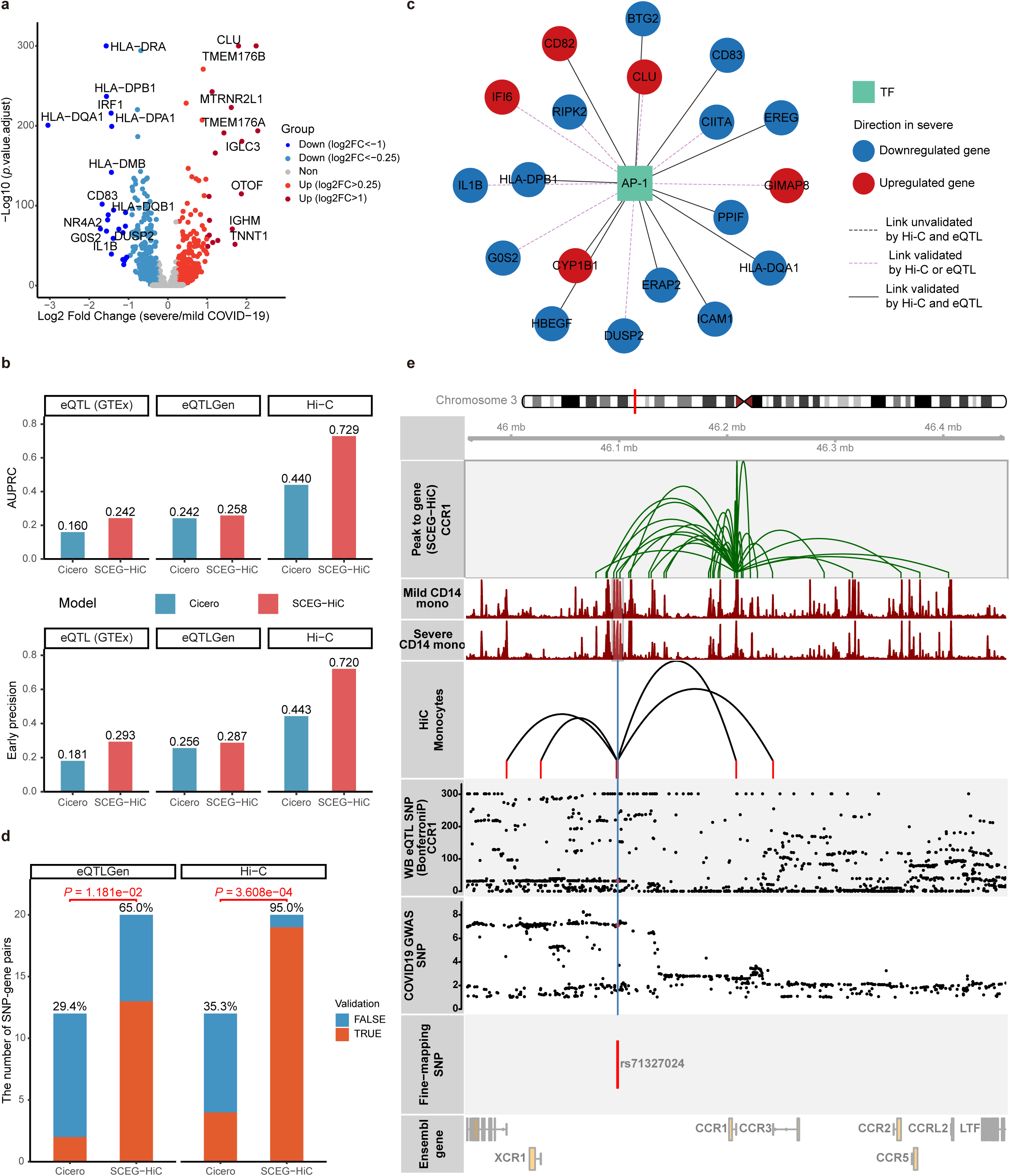
SCEG-HiC enables GRN construction and non-coding variant annotation in COVID-19 single-cell data. **a**, Volcano plot showing DEGs in scRNA-seq data between CD14+ monocytes from severe and mild COVID-19 patients. Genes with adjusted *P-*value < 0.05 and log2-FC > 0.25 are shown in red; those with log2-FC < −0.25 are shown in blue. **b**, Bar plots comparing performance between SCEG-HiC and Cicero in CD14+ monocytes, evaluated by AUPRC (top) and EP (bottom). The validation was performed using CD14+ monocyte Hi-C, and whole blood eQTL data from GTEx and eQTLGen. **c**, Subnetwork of the GRN between AP-1 TF families and target genes, inferred from TF motif enrichment analysis of SCEG-HiC-predicted enhancer-gene links. **d**, Comparison of the number of SNP-gene pairs predicted by SCEG-HiC and Cicero, with the validation fractions shown using CD14+ monocyte Hi-C and eQTLGen data. *P*-values were calculated using Fisher’s exact tests. **e**, SCEG-HiC prediction of fine-mapped SNP rs71327024 linked to *CCR1*. Integration of SCEG-HiC-predicted enhancer-gene links, scATAC-seq signals, CD14+ monocyte Hi-C interactions, whole blood eQTL SNPs, and COVID-19 GWAS SNPs around rs71327024.

We first evaluated the accuracy of predicting enhancer-DEG links using AUPRC and EP. We benchmarked against three external validation datasets: CD14+ monocyte Hi-C data from ENCODE^50^, whole blood eQTL data from eQTLGen^60^ and GTEx^51^. Across all benchmarks, SCEG-HiC consistently outperformed Cicero in both AUPRC and EP, demonstrating its superior accuracy in identifying enhancer-gene links for a specific cell type (Fig. 6b). In addition, we separately used CD14+ monocyte scATAC-seq data from severe and mild COVID-19 patients to predict enhancer-DEG links. The results showed that SCEG-HiC consistently achieved higher AUPRC and EP than Cicero in both patient groups, reinforcing its robustness across varying disease states (Supplementary Fig. 8b, c).

Next, we constructed GRNs using TFs linked to enhancer-DEG links predicted by the model, focusing on 41 most significantly DEGs in CD14+ monocytes (|log2-FC| > 1, adjusted *P* < 0.05). Using both SCEG-HiC and Cicero, we predicted enhancers associated with these DEGs and selected the top 80 enhancer-DEG links with the highest prediction scores from each model. Motif enrichment analysis was then performed on the predicted enhancer regions to identify significantly enriched TFs (Methods). Among the top 80 predicted links, SCEG-HiC identified significantly more links supported by external datasets compared to Cicero (Hi-C: *P* = 1.848 × 10^−7^; eQTLGen: *P* = 0.037; GTEx: *P* = 0.017 via one-sided Fisher’s exact test). Motif enrichment analyses based on both models revealed significant enrichment of C/EBP and AP-1 TF families (Supplementary Table 5). Previous studies have shown that AP-1 and C/EBP are associated with the immune activity of classical monocytes in COVID-19 patients^61^. After viral infection, host cells secrete interferons, activating signaling pathways such as JAK-STAT, which in turn regulate AP-1 and C/EBP TFs and further influence the expression of inflammatory genes like *IL1B*^62^. SCEG-HiC more reliably recapitulated enhancer-mediated relationships between these TFs and their target genes, as shown in GRNs inferred from predicted links (Fig. 6c, Supplementary Fig. 8d, Supplementary Table 6 and Methods).

Finally, we explored the application of SCEG-HiC in linking non-coding variants to their target genes. We retrieved 265 fine-mapped SNPs from Yu et al.^63^, which were derived from GWAS analyses comparing hospitalized COVID-19 patients with infected non-hospitalized individuals^64^. Given that these SNPs were shown to be enriched in the accessible chromatin regions of CD14+ monocytes^63^, we leveraged CD14+ monocyte scATAC-seq data to predict their target genes. SCEG-HiC and Cicero identified 20 and 17 SNP-gene links, respectively, which were validated against CD14+ monocyte Hi-C and eQTLGen data. SCEG-HiC validated 19/20 (95.0%) SNP-gene links with Hi-C data and 13/20 (65.0%) with eQTLGen data. In contrast, Cicero validated only 7/17 (35.3%) links with Hi-C data and 6/17 (29.4%) with eQTLGen data. These results demonstrated that SCEG-HiC significantly outperformed Cicero in the accuracy of predicting SNP-gene links (Hi-C: *P* = 3.608 × 10^−4^; eQTLGen: *P* = 1.181 × 10^−2^ via one-sided Fisher’s exact test; Fig. 6d).

As shown in Fig. 6e, we illustrated a potential regulatory program indicating the effect of a fine-mapped SNP, rs71327024, in the context of COVID-19. Specifically, SCEG-HiC linked this SNP to *CCR1* based on CD14+ monocyte scATAC-seq data from COVID-19 patients, an association supported by both CD14+ monocyte Hi-C and whole blood eQTL data. In contrast, Cicero linked it to *CCRL2*, a prediction lacking external validation. Besides, previous studies have confirmed that *CCR1* is a critical mediator of monocyte/macrophage polarization and tissue infiltration, which are pathogenic hallmarks of severe COVID-19^65, 66^. Multiple risk SNPs have been shown to be associated with the *CCR* gene cluster (*CCR1*, *CCR2*, *CCR3*, and *CCR5*) in monocytes/macrophages, whereas *CCRL2* exhibits no such associations^67^. We also identified another example where *CXCR6* was associated with multiple fine-mapped SNPs, which were validated by Hi-C and eQTL data (Supplementary Fig. 8e). The chemokine receptor CXCR6 and its ligand CXCL16 were ever found to play an important role in the immunopathogenesis of severe COVID-19^68^. In contrast, Cicero failed to identify any genes associated with these SNPs. A PheWAS analysis also identified several monocyte-related phenotypes associated with SNPs in the *CXCR6* region, indicating a functional role of *CXCR6* in monocytes^69, 70^. These results underscored SCEG-HiC’s superiority in identifying biologically plausible SNP-gene associations.

## Discussions

In this study, we presented SCEG-HiC, a machine learning framework that integrates single-cell omics correlations with prior chromatin conformation information to predict enhancer-gene links. SCEG-HiC is applicable to both paired scATAC/RNA-seq data and scATAC-seq data alone. Using ten single-cell multi-omics datasets from both humans and mice, we systematically benchmarked SCEG-HiC across three scenarios: (i) comparison with methods for paired multi-omics in human datasets, (ii) comparison with methods for scATAC-seq-only in human datasets, and (iii) comparison with methods for paired multi-omics in mouse datasets. SCEG-HiC consistently demonstrated high accuracy and robust performance in predicting enhancer-gene links across diverse tissues and cell types, validated using available cell type-specific Hi-C and eQTL data. While the ABC model and SCEG-HiC exhibited distinct advantages in different evaluation metrics, SCEG-HiC demonstrated a significant advantage in capturing concordance between predicted cumulative enhancer activity and gene expression. Finally, leveraging scATAC-seq data from COVID-19 patient PBMCs, we demonstrated the utility of SCEG-HiC in constructing cell type-specific GRNs by linking peaks with TFs, and prioritizing associations between non-coding variants and their potential target genes.

The key innovation of SCEG-HiC lies in its effective incorporation of prior knowledge to predict enhancer-gene links. Current methods for inferring gene regulatory relationships from single-cell omics data, such as Signac^29^, FigR^30^, Cicero^33^, SCENIC+^31^ and DIRECT-NET^32^, primarily rely on inherent statistical correlations of the data (e.g., gene expression, chromatin accessibility). However, these purely data-driven approaches have several limitations, particularly the assumption that statistical correlations directly reflect causal relationships. In practice, correlation does not necessarily imply causation, and this assumption can lead to false discoveries, especially in scenarios involving co-regulation, indirect regulation, or the combinatorial influence of complex factors such as TFs, chromatin structure, and cellular states^71, 72^. Integrating prior knowledge as a structural constraint helps increase confidence in biologically meaningful connections, suppress spurious correlations, and improve both model generalizability and interpretability. Specifically, SCEG-HiC incorporates bulk average Hi-C data as prior information about enhancer-promoter spatial contacts—a fundamental biophysical feature—alongside signals from scATAC-seq and scRNA-seq data. This integration enhances the accuracy and biological plausibility of regulatory inference. Importantly, SCEG-HiC incorporates the prior information as a tunable penalty term in sparse graphical model estimation, balancing model generalizability and predictive performance.

SCEG-HiC shows strong potential for broad applications in various biological research areas. On the one hand, combining with TF binding site prediction and TF motif enrichment analysis enables systematical construction of enhancer-mediated GRNs, offering insights into cell type-specific regulatory programs underlying diverse biological processes. On the other hand, SCEG-HiC facilitates functional annotation of non-coding genetic variants from GWAS by accurately mapping them to their putative target genes. This capacity illuminates molecular mechanisms underlying complex traits, accelerates identification of causal genes and prioritization of candidate therapeutic targets.

While SCEG-HiC demonstrates robust performance and broad application potential, several limitations warrant consideration. Firstly, its dependence on bulk average Hi-C data as prior knowledge limits applicability to species with scarce chromatin interaction resources. Secondly, while the linear structure of wglasso provides good interpretability, it may overlook complex nonlinear regulatory mechanisms. Additionally, SCEG-HiC constructs static regulatory links that do not explicitly account for dynamic regulatory changes, necessitating further extensions to capture time-dependent processes such as cell fate transitions or disease progression.

In conclusion, SCEG-HiC establishes a computational framework that integrates prior Hi-C information to infer enhancer-gene links from single-cell omics data. It offers powerful computational tool for constructing transcriptional regulatory maps, and advancing our understanding of the molecular mechanisms underlying complex traits and major diseases.

## Methods

### Data preprocessing of single-cell data

Aggregation and single-cell retention approaches are developed separately for single-cell omics data. We employed an aggregation method analogous to DIRECT-NET, which involves constructing a low-dimensional *k*-nearest neighbor (KNN) graph (default *k* = 50) from sparse single-cell omics data and subsequently averaging signals from similar cells^32^. For paired scATAC/RNA-seq data, the KNN graph is constructed using the WNN approach implemented in the Seurat v4 package, which computes a joint neighbor graph based on a weighted integration of chromatin accessibility and gene expression profiles^45^. For scATAC-seq data alone, the KNN graph is built through singular value decomposition (SVD) on a transformed peak count matrix using term frequency-inverse document frequency (TF-IDF)^29^. The aggregation process follows a modified DIRECT-NET approach, selecting cells with minimal KNN overlap (default 0.8). Finally, the aggregated chromatin accessibility and gene expression data are normalized using scaling factors estimated by DESeq2^73^. The single-cell retention approach requires normalization of both scATAC-seq and scRNA-seq data. We employed the SCTransform method in Seurat to normalize gene expression data, correcting for sequencing depth differences^74^. Chromatin accessibility data is normalized using TF-IDF method in Signac, adjusting for sequencing depth and peak accessibility biases^29^.

In this study, we evaluated the impact of six different data preprocessing methods on model accuracy, considering three factors including cell aggregation, scATAC-seq binarization and cell type scope: (i) aggregation using raw scATAC-seq data; (ii) aggregation using binarized scATAC-seq data; (iii) single-cell retention using TF-IDF normalized scATAC-seq data for all cell types; (iv) single-cell retention using binarized scATAC-seq data for all cell types; (v) single-cell retention using TF-IDF normalized scATAC-seq data for a specified cell type; and (vi) single-cell retention using binarized scATAC-seq data for a specified cell type.

### Constructing and normalizing contact matrix of bulk average Hi-C

To obtain the bulk averaged Hi-C data for constructing the enhancer-gene contact matrix, we adopted a similar method to the ABC model^23^. First, we extracted rows corresponding to the TSS of each gene from the standardized Hi-C matrix for each cell type (KR or SCALE normalization, at 5 kb resolution)^75^. Considering that the strength of intra-chromosomal interactions follows a power law distribution, we corrected for differences in the power-law decay of Hi-C signals across different cell types^76^. This is achieved by adjusting the signal based on the cell-type specific power law gamma parameter, using the formula:

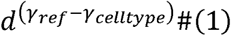

Where *γ_ref_* is the reference gamma parameter, ensuring a consistent decay pattern across all cell types that matches the reference. To estimate *γ_celltype_*, we aggregated Hi-C interactions within 1 Mb of all gene promoters and performed a linear regression in log-log space, where the regression slope corresponded to *γ_celltype_*. After scaling, each Hi-C profile was normalized so that the total signal summed to one, ensuring equal weighting across all cell types. Finally, we averaged the normalized profiles across all cell types to construct a consensus Hi-C interaction landscape anchored at gene TSSs.

We extracted a contact matrix from the bulk average Hi-C data using the TSS of genes and the midpoints of all candidate enhancers. The coordinates were divided by 5 kb (the Hi-C resolution) and then floor-rounded using R’s floor function. Given the low contact frequency of the resulting matrix, we standardized the data to the [0, 1] range and used it as a penalty matrix for subsequent wglasso modeling, with larger contact frequencies assigned smaller penalties. Four standardization methods were evaluated: (i) binarization, where contact frequencies are assigned to 0 (contact present) or 1 (contact absent); (ii) min-max normalization, where contact frequencies are scaled to the [0, 1] range by subtracting each value from their maximum and dividing by their range (max-min); (iii) -log transformation, where contact frequencies are -log-transformed and then min-max normalized; (iv) rank score, where contact frequencies are ranked in descending order and scaled to the [0, 1] range by relative ranks.

### Construction of a predictive enhancer-gene links model

SCEG-HiC employs the wglasso method to predict enhancer-gene links, effectively handling the high dimensionality of single-cell datasets, where the number of potential element pairs far exceeds the cell number^36^. Moreover, wglasso integrates multi-level biological information, enhancing model accuracy and robustness. Like the glasso, wglasso estimates the inverse covariance matrix to compute partial correlation coefficients, thereby removing indirect links^77^. The key difference is that while glasso applies a single penalty parameter, wglasso leverages prior information to specify different amounts of penalties to different element pairs, enabling adaptive shrinkage of partial correlations for improved accuracy. For each gene, wglasso maximizes the following objective function:

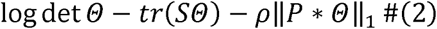

where *Θ* is the inverse covariance matrix to be estimated, *S* is the observed covariance matrix, *P* is the prior information matrix (*P ∈* [0,1]), p is the penalty parameter, and * represents element-wise multiplication. With scATAC-seq data alone, the covariance matrix is computed based on the chromatin accessibility of promoters (peaks within ± 1 kb from the TSS) and candidate enhancers (peaks within ± 250 kb from the TSS). When paired scATAC/RNA-seq data are available, the covariance matrix is computed using gene expression and enhancer accessibility. The standardized contact matrix derived from bulk average Hi-C data serves as the prior information matrix *P*, where larger values indicate stronger penalties and weaker prior support for corresponding enhancer-gene pairs. To ensure numerical stability during optimization, a small constant (1X 10^−4^) is added to the diagonal of *P*. The optimal penalty parameter is determined by cross-validation using the BIC, which tends to select highly sparse networks and performs well in scale-free network simulations^40^. The BIC for the wglasso model takes the form

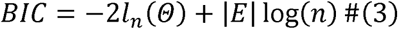

where |*E*| denotes the number of edges, *n* is the sample size, and *l_n_* is the log likelihood function simplified from

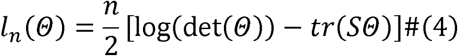

After estimating the inverse covariance matrix, partial correlation coefficients are calculated to infer enhancer-gene links, with a default threshold of 0.01. The partial correlation coefficient *R* is calculated as:

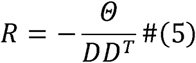

where *D* is a diagonal matrix composed of the square roots of the diagonal elements of *θ*.

### Preprocessing paired scRNA-seq and scATAC-seq datasets

Single-cell paired multi-omics data were processed using Seurat (scRNA-seq) and Signac (scATAC-seq). QC was applied to both the RNA and ATAC data, assessing RNA counts, ATAC fragment counts, nucleosome signal, TSS enrichment, and optionally blacklist fraction. QC thresholds were defined based on the distribution of these metrics within each dataset, as detailed in Supplementary Table 7. For scRNA-seq data, RNA counts were normalized using Seurat’s SCTransform function, followed by dimensionality reduction with RunPCA. For scATAC-seq data, MACS2 was used via Signac’s CallPeaks function to identify peaks from fragment files^59^. After filtering out peaks on non-standard chromosomes and overlapping with blacklisted genomic regions, a peak-by-cell matrix was generated using Signac’s FeatureMatrix function for downstream analyses. ATAC peaks were normalized using FindTopFeatures and RunTFIDF in Signac, and dimensionality reduction was performed using RunSVD. If the dataset contained multiple samples, batch effects based on processing date were corrected using the Harmony method with the RunHarmony function^78^. The WNN graph was constructed using latent semantic indexing (LSI) dimensions 2-40 for scATAC-seq (excluding the first singular component due to its strong correlation with technical factors like sequencing depth) and PCA dimensions 1-50 for scRNA-seq. Finally, high-resolution clustering was performed using the Louvain algorithm on the joint WNN graph via Seurat’s FindNeighbors and FindClusters functions.

### Cell type annotation

Cell type annotation was provided by the original studies for mouse skin. For other datasets including human skin stromal^41^, fetal retinal^42^, brain gray matter^43^, and developing cerebral cortex^44^, as well as mouse cerebral cortex^55^ and thymic epithelial cells^56^, we performed clustering and annotation using previously reported marker genes. Notably, for the adult mouse cerebral cortex, we carefully distinguished between subtypes of excitatory and inhibitory neurons. For human PBMCs, cell annotation was further examined using the PBMC reference dataset from Hao et al.^45^ and Seurat’s label transfer functions FindTransferAnchors and TransferData. Embryonic mouse brain cells were annotated based on the TRIPOD model’s cell annotation tutorial (https://github.com/yuchaojiang/TRIPOD/blob/main/scripts/10x_e18/e18_01_create_seurat_object.R)^79^. Mouse liver cells were clustered and annotated according to Carmen et al. ’s annotation results, with clusters renamed based on the cell type showing the greatest overlap^57^.

### Existing methods for paired scATAC-seq/scRNA-seq data

We compared SCEG-HiC with several existing methods for paired scATAC/scRNA data, including two classical correlation-based approaches (Pearson and Spearman), as well as five computational tools. (i) CellOracle (v0.14.0), which integrates Cicero results with TSS peak information to infer peak-gene links from scATAC-seq data^47^; (i) DIRECT-NET (v0.0.1), which applies eXtreme Gradient Boosting (XGBoost) to identify high-confidence functional CRE-gene links from parallel scATAC/RNA-seq data^32^; (iii) SCENIC+ (v1.0.1), which utilizes gradient boosting machine (GBM) regression to compute CRE-gene importance scores from scATAC/RNA-seq data^31^; (iv) FigR (v0.1.0), which employs Spearman correlation coefficients to connect distal CREs to genes from paired scATAC/RNA-seq data^30^; (v) Signac (v1.12.0), which applies linear regression-based methods to compute linkage scores for peak-gene pairs from scATAC/RNA-seq data^29^. For all models, promoters are defined as regions within ± 1 kb from the TSS, while candidate enhancers are defined as regions within ± 250 kb from the TSS, excluding promoter regions. A common set of marker genes were selected for each cell type, and all resulting links were retained for benchmarking purposes.

For two classical correlation-based approaches, Pearson and Spearman correlations were computed using R’s cor function with method = “pearson” or method = “spearman”, respectively, applied to gene expression and candidate enhancer accessibility after aggregating the data.

For CellOracle (v0.14.0), which builds on Cicero (v1.3.9) for inferring regulatory links, we followed the official tutorial (https://cole-trapnell-lab.github.io/cicero-release/docs_m3/). Since Cicero only supports for GRCh37 and GRCm38 genome assemblies, we used the liftOver function from the rtracklayer package to convert GRCh38 coordinates to GRCh37 when necessary^80^. We then binarized scATAC-seq data and used the new_cell_data_set function to create cell_data_set (CDS) objects. Next, Cicero was run with default parameters for data preprocessing, including detect_genes, estimate_size_factors, preprocess_cds, reduce_dimension and make_cicero_cds. Finally, the run_cicero function was used to identify peak-peak co-accessibility scores. Finally, CellOracle incorporated Cicero’s results with TSS peak information and filtered peaks with weak connections to TSS peaks (coaccess > 0).

For DIRECT-NET (v0.0.1), we followed the official example (https://htmlpreview.github.io/?https://github.com/zhanglhbioinfor/DIRECT-NET/blob/main/tutorial/demo_DIRECTNET_PBMC.html). We modified the code to unify promoters and enhancers, then ran the Run_DIRECT_NET function to predict the links between CREs and target genes. The parameters for the function were k_neigh = 50, atacbinary = TRUE, max_overlap = 0.5, and size_factor_normalize = TRUE.

For SCENIC+ (v1.0.1), we first imputed accessibility count matrices using pyCisTopic, inferring 50 topics. TF binding was inferred via the CisTarget and DEM motif matcher algorithms, utilizing the CisTargetDB motif collection. The scRNA-seq Seurat object was converted to an AnnData object via the seurat2anndata function. All data were then integrated into a scenicplus object. To predict peak-gene links, we used the get_search_space and calculate_regions_to_genes_relationships functions. The parameters for the get_search_space function were biomart_host = ‘http://www.ensembl.org’, upstream = [1000, 250000], downstream = [1000, 250000], and extend_tss = [1000, 1000].

For FigR (v0.1.0), we followed the official tutorial (https://buenrostrolab.github.io/FigR/). First, we used the SEfromSignac function to create a SummarizedExperiment object containing peak read counts derived from the Seurat object. Next, we combined it with the paired sparseMatrix of scRNA-seq gene counts, and applied the runGenePeakcorr function to determine peak-gene links. The parameters for the function were genome = “hg38” or “mm10”, and p.cut = NULL to retain all tested links.

For Signac (v1.12.0), we used the MatchRegionStats function to generate a background distribution for each peak by selecting 200 random peaks with matched GC content, accessibility, and sequence length. Then, the LinkPeaks function was applied to compute the correlation coefficient between gene expression and CRE accessibility within a given distance from the gene TSS. This observed correlation was compared against an expected correlation derived from the background to compute *z*-scores and *P*-values. The parameters for the function were method = “pearson”, distance = 250000, pvalue_cutoff = 1.01, and score_cutoff = 0 to retain all tested links.

### Existing methods for scATAC-seq data alone

We compared SCEG-HiC with several existing methods for scATAC-seq data alone, including two statistical approaches: Pearson correlation and Chi2+FDR, as well as three computational tools. (i) Cicero (v1.3.9), which uses the glasso model to infer co-accessibility relationships between CREs from scATAC-seq data alone^33^; (ii) scEChIA (v0.1.0), which utilizes signal imputation and L1 regularization to infer chromatin interaction links from scATAC-seq data^54^; (iii) DIRECT-NET (v0.0.1), which applies XGBoost to identify high-confidence functional gene-CRE links from scATAC-seq data alone^32^. All methods followed the same promoter and enhancer definitions as previously described to ensure consistency across evaluations.

For Pearson correlation, Cicero (v1.3.9), and DIRECT-NET (v0.0.1), we followed the same analysis pipeline as described above, except that scATAC-seq data alone served as the input.

For the Chi2+FDR method, scATAC-seq data were first binarized, and 2 × 2 contingency tables were constructed for each promoter-enhancer pair. R’s chisq.test function was used to calculate *P*-values, which were corrected for multiple hypothesis testing using the Benjamini-Hochberg FDR procedure.

For scEChIA (v0.1.0), we followed the official tutorial (http://reggen.iiitd.edu.in:1207/scEChIA/). We used the Hi-C data (25 kb resolution) from GM12878 and IMR90 cell lines provided in scEChIA. We applied the rhomatAvg function to compute the average of these two Hi-C datasets. Interaction predictions were then performed using the Interaction_Prediction_1 function, with parameters including chrinfo, rhomatrix, and chromSize. Like Cicero, scEChIA relies on the GRCh37 genome assembly, requiring the use of the liftOver function to convert GRCh38 coordinates to GRCh37.

### ABC model

Additionally, we compare SCEG-HiC with the ABC method for bulk ATAC-seq, which integrates chromatin accessibility and 3D genome contact frequency to predict enhancer-gene links^23^. To maintain consistency, ABC analysis employed the same promoter and enhancer definitions as described above.

For ABC, we followed the official tutorial (https://github.com/broadinstitute/ABC-Enhancer-Gene-Prediction). To generate the required input files (EnhancerList.txt and GeneList.txt), we first calculated the average gene expression and chromatin accessibility across each cell type using the AverageExpression function from the Seurat package. Candidate peaks were then annotated using the annotatePeak function from the ChIPseeker package with the parameter tssRegion = c (−1000, 1000)^81^. Peaks are categorized as “Distal Intergenic”, “Promoter”, or “Others” based on their genomic location. The resulting data were then formatted to meet the input requirements of the ABC model. Subsequently, ABC scores were computed using the predict.py script with parameters: --window 250000, --expression_cutoff 0.0, --tss_slop 1000, -- promoter_activity_quantile_cutoff 0.0, --cellType average, and --threshold 0.0. All other necessary input files were prepared following the official documentation.

### Validation with Hi-C and eQTL data

To evaluate the accuracy of predicted enhancer-gene links, we used published cell type-specific Hi-C profiles (Supplementary Table 8). We used Juicer Tools to process raw sequencing data into a list of Hi-C contacts (.hic files)^82^. The chromatin interaction scores were normalized using the SCALE or KR methods and extracted in text format at 5 kb resolution. An enhancer-gene link is considered as a true positive (TP) if: (i) the gene TSS is within a bait fragment, (ii) the midpoint of the peak is within an other-end fragment, and (iii) a high interaction score is observed between the bait and other-end fragment in Hi-C data, determined by a threshold to select top-enriched chromatin interactions. Additionally, we used eQTL data from matched tissues in GTEx (v8) to assess the accuracy of enhancer-gene links (Supplementary Table 8). Specifically, we used cis-eQTL data, which link genetic variants to the expression of nearby genes. An enhancer-gene link is considered as a TP if (i) the peak overlaps with an eQTL locus, and (ii) the locus is associated with the target gene’s expression.

### Evaluation metrics

To derive a common gene set for benchmarking, we first identified 3,000 highly variable genes for each dataset. We then used the presto package to select marker genes for each cell type with AUROC > 0.5^83^. The evaluation metrics for comparison included the AUPRC, AUPRC ratio, EP, and EPR^49^. AUPRC was computed using the pr.curve function from the PRROC package, and EP was defined as the fraction of TPs among the top-*k* predicted positives, where k represents the total number of TPs. Both the AUPRC ratio and EPR were calculated by dividing the AUPRC and EP values by the precision of a random predictor, which is determined by the total fraction of TPs.

### Correlations between cumulative enhancer activity and gene expression

To calculate cumulative enhancer activity, we first normalized the peak counts by the total number of unique fragments in each cell’s peaks. After normalization, the cumulative enhancer activity for each gene is defined as the sum of the normalized counts from all significantly associated peaks, resulting in a cell *X* cumulative enhancer activity matrix. The thresholds for significant peaks were model-specific (ABC: 0.02, SCEG-HiC: 0.001). We then computed the Spearman correlations between cumulative enhancer activity and gene expression values across cells for each gene.

### Selection and DEG analysis of COVID-19 scRNA-seq data

The scRNA-seq dataset was processed and annotated by Wilk et al. following the Seurat workflow^58^. Since our primary focus was comparing PBMC samples from mild and severe COVID-19 patients, we excluded samples derived from red blood cell-lysed whole blood and removed individuals annotated as “healthy” or “convalescent”. To compare mild and severe COVID-19 patients, differential expression tests were performed for each cell type using Seurat’s FindMarkers function with the Wilcoxon rank-sum test. Genes were considered significantly differentially expressed if they met all the following criteria: (i) absolute log2-FC 2: 0.25, (ii) Bonferroni-corrected *P*-value < 0.05, and (iii) expression in at least 10% of cells in either group.

### Processing and reference-based annotation of COVID-19 scATAC-seq data

We processed each COVID-19 scATAC-seq sample using Signac^29^. Given that only fragment files were available, we first generated the count matrix with the FeatureMatrix function and then identified peaks using the CallPeaks function with MACS2^59^. For each sample, we applied QC filters based on TSS enrichment, nucleosome signal, and log10 fragment count per cell. QC thresholds were determined individually for each sample to account for variations in sequencing depth and sample quality, as detailed in Supplementary Table 7.

For cell type annotation, we used a publicly available paired multi-omics dataset of healthy PBMCs from 10x Genomics (https://support.10xgenomics.com/single-cell-multiome-atac-gex/datasets/1.0.0/pbmc_granulocyte_sorted_10k), which is suitable for integration with both scRNA-seq and scATAC-seq data. The RNA and ATAC count matrices were filtered to retain cells with RNA reads between 1,000 and 15,000, ATAC fragments between 1,000 and 100,000, nucleosome signal less than 2, TSS enrichment greater than 1, and less than 20% mitochondrial gene expression. Peak calling on ATAC fragment files was performed using MACS2 via Signac’s CallPeaks function, and the peak count matrix was generated with the FeatureMatrix function. Cell type annotations for the multi-omics dataset were performed using the Azimuth tool from Seurat v4, following the same procedure as for the scRNA-seq data^45^. To reduce the dimensionality of the peak matrix, we performed LSI using Signac’s RunTFIDF and RunSVD functions. This approach establishes an ATAC-based dimensionality space integrated with RNA-based automated cell type annotations from the reference dataset. Finally, we projected each COVID-19 scATAC-seq sample into the dimensionality space of the paired multi-omics dataset, transferring cell type annotations to our scATAC-seq data using Seurat’s FindAnchors and TransferData functions.

To integrate the peak count matrices across samples, we used Signac’s merge function to recalculate peak counts based on a unified set of common peaks. Dimensionality reduction was then performed on the integrated dataset, followed by batch effect correction across samples using the RunHarmony function. Finally, cell clusters were identified using Seurat’s FindClusters function on the batch-corrected low-dimensional embeddings, and each cluster was relabeled according to the most enriched known cell type.

### TF motif enrichment analysis

TF motif enrichment analysis was performed using motif position weight matrices (PWM) from the JASPAR2020 database^84^. Core TF motifs were retrieved using the getMatrixSet function from the TFBSTools package^85^ with parameters species = “Homo sapiens”, collection = “CORE”, and all_versions = FALSE, resulting in 633 core TF motifs. Genomic region statistics—including GC content, sequence length, and dinucleotide frequencies—were computed using the RegionStats function. Motif annotations were then added to the object using AddMotifs. Finally, we selected relevant peaks and performed motif enrichment analysis within these regions with the FindMotifs function. Motifs with a fold enrichment greater than 2 and an adjusted *P*-value less than 0.05 were considered significant.

### Construction of GRNs

To construct GRNs, we used the matchMotifs function from the motifmatchr package to identify TFs associated with peak regions, leveraging motif data from the JASPAR2020 database^86^. Only TF motifs showing significant enrichment in these regions were considered. We integrated these TF-peak links with peak-gene links derived from SCEG-HiC to establish TF-peak-gene relationships. In this framework, peaks serve as intermediaries linking TFs to their putative target genes, forming a GRN structure.

## Data availability

All datasets used in this study are publicly available without restrictions. The paired scATAC/RNA-seq datasets were obtained from the 10X Genomics website and the Gene Expression Omnibus (GEO) database: PBMC (https://www.10xgenomics.com/datasets/pbmc-from-a-healthy-donor-granulocytes-removed-through-cell-sorting-10-k-1-standard-1-0-0), skin stromal (<u>GSE195452</u>), fetal retina (<u>GSE246169</u>), brain gray matter (<u>GSE193240</u>), developing cerebral cortex (<u>GSE162170</u>), mouse embryonic brain cells at day 18 (https://www.10xgenomics.com/datasets/fresh-embryonic-e-18-mouse-brain-5-k-1-standard-2-0-0), mouse skin (<u>GSE140203</u>), adult mouse cerebral cortex (<u>GSE126074</u>), thymic epithelial (<u>GSE236075</u>), and mouse liver (<u>GSE218468</u>). Cell type-specific Hi-C data used for benchmarking, as well as mouse Hi-C datasets used to generate the average Hi-C, were obtained from the ENCODE project and GEO database: CD4+ T (<u>ENCFF958DWQ</u>), CD8+ T (<u>ENCFF009ONH</u>), CD14+ monocyte (<u>ENCFF185AYZ</u>), brain midfrontal cortex (<u>ENCSR165UJN</u>), skin fibroblasts (<u>GSM5266524</u>), retinal (<u>GSE229683</u>), glutamatergic neurons (<u>GSE228118</u>), mouse embryonic forebrain (<u>GSE201186</u>), mouse cortex(<u>GSE34587</u>), skin epidermis/suprabasal (<u>GSE197024</u>), mTEC (<u>GSE180933</u>), liver (<u>GSE65126</u>), mouse embryonic stem cell (<u>ENCFF409ZRS</u>, <u>ENCFF584EDJ</u>), CH12F3 (<u>ENCFF909ODS</u>), mature B cell (<u>ENCFF026SNO</u>), CH12.LX (<u>ENCFF076GYR</u>), and epithelium/fiber (<u>GSE243851</u>). The eQTL data used for benchmarking were mainly from the GTEx eQTL summary statistics (v8) (https://www.gtexportal.org/home/datasets), retinal eQTL summary statistics (https://www-huge.uni-regensburg.de/databases.html), and eQTLGen summary statistics (release 2019-12-11) (https://www.eqtlgen.org). The COVID-19 PBMC scATAC-seq and scRNA-seq data were downloaded from GEO (<u>GSE174072</u>). The human bulk average Hi-C data were obtained from the ENCODE project (<u>ENCFF134PUN</u>). Details of the data sources are provided in Supplementary Table 2, 4, 8. The processed single-cell data used in this study, along with the mouse bulk average Hi-C data, are available via Zenodo: https://zenodo.org/record/14849886.

## Code availability

SCEG-HiC is available as an R package at https://github.com/wuwei77lx/SCEGHiC. A GitHub repository containing all original code used for running and evaluating existing models is available at https://github.com/wuwei77lx/compare_model.

## Acknowledgements

This work was supported by the National Key R&D Program of China (2021YFA1100501), the National Natural Science Foundation of China (32370690) and the Major Project of Guangzhou National Laboratory (GZNL2024A01003). We thank Prof. Sijia Wang for helpful discussions on this work.

## Author Contributions

X.L. and Z.W. designed the research. X.L. performed data analysis. X.L., Y.M. and D.H. built and tested the software. X.L. wrote the manuscript. Y.M., D.H., Y.L., W.Z. and Z.W. revised the manuscript. Z.W. supervised the study.

## References

1. Casamassimi, A. & Ciccodicola, A. Transcriptional Regulation: Molecules, Involved Mechanisms, and Misregulation. Int. J. Mol. Sci. 20, 1281 (2019).

2. Lenhard, B., Sandelin, A. & Carninci, P. Metazoan promoters: emerging characteristics and insights into transcriptional regulation. Nat. Rev. Genet. 13, 233-245 (2012).

3. Pennacchio, L.A., Bickmore, W., Dean, A., Nobrega, M.A. & Bejerano, G. Enhancers: five essential questions. Nat. Rev. Genet. 14, 288–295 (2013).

4. Claringbould, A. & Zaugg, J.B. Enhancers in disease: molecular basis and emerging treatment strategies. Trends Mol. Med. 27, 1060–1073 (2021).

5. Sanyal, A., Lajoie, B.R., Jain, G. & Dekker, J. The long-range interaction landscape of gene promoters. Nature 489, 109–113 (2012).

6. Zhang, Y. et al. Chromatin connectivity maps reveal dynamic promoter-enhancer long-range associations. Nature 504, 306–310 (2013).

7. Dixon, J.R. et al. Chromatin architecture reorganization during stem cell differentiation. Nature 518, 331–336 (2015).

8. van Arensbergen, J., van Steensel, B. & Bussemaker, H.J. In search of the determinants of enhancer-promoter interaction specificity. Trends Cell Biol. 24, 695–702 (2014).

9. Dekker, J., Rippe, K., Dekker, M. & Kleckner, N. Capturing chromosome conformation. Science 295, 1306–1311 (2002).

10. van Berkum, N.L. et al. Hi-C: a method to study the three-dimensional architecture of genomes. J. Vis. Exp. 39, 1869 (2010).

11. GTEx Consortium. Genetic effects on gene expression across human tissues. Nature 550, 204–213 (2017).

12. Gasperini, M. et al. A Genome-wide Framework for Mapping Gene Regulation via Cellular Genetic Screens. Cell 176, 377–390.e319 (2019).

13. Moore, J.E., Pratt, H.E., Purcaro, M.J. & Weng, Z. A curated benchmark of enhancer-gene interactions for evaluating enhancer-target gene prediction methods. Genome Biol. 21, 17 (2020).

14. Hariprakash, J.M. & Ferrari, F. Computational Biology Solutions to Identify Enhancers-target Gene Pairs. Comput. Struct. Biotechnol. J. 17, 821–831 (2019).

15. Xu, H., Zhang, S., Yi, X., Plewczynski, D. & Li, M.J. Exploring 3D chromatin contacts in gene regulation: The evolution of approaches for the identification of functional enhancer-promoter interaction. Comput. Struct. Biotechnol. J. 18, 558–570 (2020).

16. Ernst, J. et al. Mapping and analysis of chromatin state dynamics in nine human cell types. Nature 473, 43–49 (2011).

17. Thurman, R.E. et al. The accessible chromatin landscape of the human genome. Nature 489, 75–82 (2012).

18. He, B., Chen, C., Teng, L. & Tan, K. Global view of enhancer-promoter interactome in human cells. Proc. Natl. Acad. Sci. U. S. A. 111, E2191–E2199 (2014).

19. Whalen, S., Truty, R.M. & Pollard, K.S. Enhancer-promoter interactions are encoded by complex genomic signatures on looping chromatin. Nat. Genet. 48, 488–496 (2016).

20. Hafez, D. et al. McEnhancer: predicting gene expression via semi-supervised assignment of enhancers to target genes. Genome Biol. 18, 199 (2017).

21. Fishilevich, S. et al. GeneHancer: genome-wide integration of enhancers and target genes in GeneCards. Database (Oxford) 2017, bax028 (2017).

22. Zhu, Y. et al. Constructing 3D interaction maps from 1D epigenomes. Nat. Commun. 7, 10812 (2016).

23. Fulco, C.P. et al. Activity-by-contact model of enhancer-promoter regulation from thousands of CRISPR perturbations. Nat. Genet. 51, 1664-1669 (2019).

24. Tang, F. et al. mRNA-Seq whole-transcriptome analysis of a single cell. Nat. Methods 6, 377–382 (2009).

25. Buenrostro, J.D., Giresi, P.G., Zaba, L.C., Chang, H.Y. & Greenleaf, W.J. Transposition of native chromatin for fast and sensitive epigenomic profiling of open chromatin, DNA-binding proteins and nucleosome position. Nat. Methods 10, 1213–1218 (2013).

26. Ma, S. et al. Chromatin Potential Identified by Shared Single-Cell Profiling of RNA and Chromatin. Cell 183, 1103–1116.e1120 (2020).

27. Stoeckius, M. et al. Simultaneous epitope and transcriptome measurement in single cells. Nat. Methods 14, 865–868 (2017).

28. Kim, D. et al. Gene regulatory network reconstruction: harnessing the power of single-cell multi-omic data. NPJ Syst. Biol. Appl. 9, 51 (2023).

29. Stuart, T., Srivastava, A., Madad, S., Lareau, C.A. & Satija, R. Single-cell chromatin state analysis with Signac. Nat. Methods 18, 1333–1341 (2021).

30. Kartha, V.K. et al. Functional inference of gene regulation using single-cell multi-omics. Cell Genom. 2, 100166 (2022).

31. Bravo González-Blas, C., et al. SCENIC+: single-cell multiomic inference of enhancers and gene regulatory networks. Nat. Methods 20, 1355–1367 (2023).

32. Zhang, L., Zhang, J. & Nie, Q. DIRECT-NET: An efficient method to discover cis-regulatory elements and construct regulatory networks from single-cell multiomics data. Sci. Adv. 8, eabl7393 (2022).

33. Pliner, H.A. et al. Cicero Predicts cis-Regulatory DNA Interactions from Single-Cell Chromatin Accessibility Data. Mol. Cell 71, 858–871.e858 (2018).

34. Nagano, T. et al. Single-cell Hi-C reveals cell-to-cell variability in chromosome structure. Nature 502, 59–64 (2013).

35. Dautle, M.A. & Chen, Y. Single-Cell Hi-C Technologies and Computational Data Analysis. Adv. Sci. 12, e2412232 (2025).

36. Li, Y. & Jackson, S.A. Gene Network Reconstruction by Integration of Prior Biological Knowledge. G3 Genes|Genomes|Genet. 5, 1075–1079 (2015).

37. Luecken, M.D. & Theis, F.J. Current best practices in single-cell RNA-seq analysis: a tutorial. Mol. Syst. Biol. 15, e8746 (2019).

38. Yan, F., Powell, D.R., Curtis, D.J. & Wong, N.C. From reads to insight: a hitchhiker’s guide to ATAC-seq data analysis. Genome Biol. 21, 22 (2020).

39. Tang, Z. et al. CTCF-Mediated Human 3D Genome Architecture Reveals Chromatin Topology for Transcription. Cell 163, 1611–1627 (2015).

40. Foygel, R. & Drton, M. Extended Bayesian Information Criteria for Gaussian Graphical Models. Adv. Neural Inf. Process. Syst. 23, 604–612 (2010).

41. Gur, C. et al. LGR5 expressing skin fibroblasts define a major cellular hub perturbed in scleroderma. Cell 185, 1373–1388.e1320 (2022).

42. Wohlschlegel, J. et al. ASCL1 induces neurogenesis in human Müller glia. Stem Cell Rep. 18, 2400–2417 (2023).

43. Meijer, M. et al. Epigenomic priming of immune genes implicates oligodendroglia in multiple sclerosis susceptibility. Neuron 110, 1193–1210.e1113 (2022).

44. Trevino, A.E. et al. Chromatin and gene-regulatory dynamics of the developing human cerebral cortex at single-cell resolution. Cell 184, 5053–5069.e5023 (2021).

45. Hao, Y. et al. Integrated analysis of multimodal single-cell data. Cell 184, 3573–3587.e3529 (2021).

46. Nasser, J. et al. Genome-wide enhancer maps link risk variants to disease genes. Nature 593, 238–243 (2021).

47. Kamimoto, K. et al. Dissecting cell identity via network inference and in silico gene perturbation. Nature 614, 742–751 (2023).

48. Davis, J.J. & Goadrich, M.H. The relationship between Precision-Recall and ROC curves. in Proceedings of the 23rd international conference on Machine learning (ICML ’06), 233–240 (Year).

49. Pratapa, A., Jalihal, A.P., Law, J.N., Bharadwaj, A. & Murali, T.M. Benchmarking algorithms for gene regulatory network inference from single-cell transcriptomic data. Nat. Methods 17, 147–154 (2020).

50. ENCODE Project Consortium. An integrated encyclopedia of DNA elements in the human genome. Nature 489, 57–74 (2012).

51. GTEx Consortium. The GTEx Consortium atlas of genetic regulatory effects across human tissues. Science 369, 1318–1330 (2020).

52. Hinton, H.J., Alessi, D.R. & Cantrell, D.A. The serine kinase phosphoinositide-dependent kinase 1 (PDK1) regulates T cell development. Nat. Immunol. 5, 539–545 (2004).

53. Park, S.G. et al. The kinase PDK1 integrates T cell antigen receptor and CD28 coreceptor signaling to induce NF-kappaB and activate T cells. Nat. Immunol. 10, 158–166 (2009).

54. Pandey, N., Omkar, C., Mishra, S. & Kumar, V. Improving Chromatin-Interaction Prediction Using Single-Cell Open-Chromatin Profiles and Making Insight Into the Cis-Regulatory Landscape of the Human Brain. Front. Genet. 12, 738194 (2021).

55. Chen, S., Lake, B.B. & Zhang, K. High-throughput sequencing of the transcriptome and chromatin accessibility in the same cell. Nat. Biotechnol. 37, 1452–1457 (2019).

56. Givony, T. et al. Thymic mimetic cells function beyond self-tolerance. Nature 622, 164–172 (2023).

57. Bravo González-Blas, C., et al. Single-cell spatial multi-omics and deep learning dissect enhancer-driven gene regulatory networks in liver zonation. Nat. Cell Biol. 26, 153–167 (2024).

58. Wilk, A.J. et al. Multi-omic profiling reveals widespread dysregulation of innate immunity and hematopoiesis in COVID-19. J. Exp. Med. 218 (2021).

59. Zhang, Y. et al. Model-based analysis of ChIP-Seq (MACS). Genome Biol. 9, R137 (2008).

60. Võsa, U. et al. Large-scale cis- and trans-eQTL analyses identify thousands of genetic loci and polygenic scores that regulate blood gene expression. Nat. Genet. 53, 1300–1310 (2021).

61. Hwang, N. et al. Single-cell sequencing of PBMC characterizes the altered transcriptomic landscape of classical monocytes in BNT162b2-induced myocarditis. Front. Immunol. 13, 979188 (2022).

62. Mishra, B. & Ivashkiv, L.B. Interferons and epigenetic mechanisms in training, priming and tolerance of monocytes and hematopoietic progenitors. Immunol. Rev. 323, 257–275 (2024).

63. Yu, F. et al. Variant to function mapping at single-cell resolution through network propagation. Nat. Biotechnol. 40, 1644–1653 (2022).

64. COVID-19 Host Genetics Initiative Mapping the human genetic architecture of COVID-19. Nature 600, 472-477 (2021).

65. Mantovani, A. et al. The chemokine system in diverse forms of macrophage activation and polarization. Trends Immunol. 25, 677–686 (2004).

66. Merad, M. & Martin, J.C. Author Correction: Pathological inflammation in patients with COVID-19: a key role for monocytes and macrophages. Nat. Rev. Immunol. 20, 448 (2020).

67. Stikker, B.S. et al. Severe COVID-19-associated variants linked to chemokine receptor gene control in monocytes and macrophages. Genome Biol. 23, 96 (2022).

68. Payne, D.J. et al. The CXCR6/CXCL16 axis links inflamm-aging to disease severity in COVID-19 patients. Preprint at bioRxiv 10.1101/2021.01.25.428125 (2021).

69. Dai, Y. et al. Association of CXCR6 with COVID-19 severity: delineating the host genetic factors in transcriptomic regulation. Hum. Genet. 140, 1313–1328 (2021).

70. Kamat, M.A. et al. PhenoScanner V2: an expanded tool for searching human genotype-phenotype associations. Bioinformatics 35, 4851–4853 (2019).

71. Larcombe, M.R., Hsu, S., Polo, J.M. & Knaupp, A.S. Indirect Mechanisms of Transcription Factor-Mediated Gene Regulation during Cell Fate Changes. Adv. Genet. 3, 2200015 (2022).

72. Cheng, Y. et al. Co-regulation of invected and engrailed by a complex array of regulatory sequences in Drosophila. Dev. Biol. 395, 131–143 (2014).

73. Anders, S. & Huber, W. Differential expression analysis for sequence count data. Genome Biol. 11, R106 (2010).

74. Stuart, T. et al. Comprehensive Integration of Single-Cell Data. Cell 177, 1888–1902.e1821 (2019).

75. Knight, P.A. & Ruiz, D. A fast algorithm for matrix balancing. IMA J. Numer. Anal. 33, 1029–1047 (2012).

76. Sarnataro, S., Chiariello, A.M., Esposito, A., Prisco, A. & Nicodemi, M. Structure of the human chromosome interaction network. PLoS One 12, e0188201 (2017).

77. Friedman, J., Hastie, T. & Tibshirani, R. Sparse inverse covariance estimation with the graphical lasso. Biostatistics 9, 432–441 (2008).

78. Korsunsky, I. et al. Fast, sensitive and accurate integration of single-cell data with Harmony. Nat. Methods 16, 1289–1296 (2019).

79. Jiang, Y. et al. Nonparametric single-cell multiomic characterization of trio relationships between transcription factors, target genes, and cis-regulatory regions. Cell Syst. 13, 737–751.e734 (2022).

80. Lawrence, M., Gentleman, R. & Carey, V. rtracklayer: an R package for interfacing with genome browsers. Bioinformatics 25, 1841–1842 (2009).

81. Yu, G., Wang, L.G. & He, Q.Y. ChIPseeker: an R/Bioconductor package for ChIP peak annotation, comparison and visualization. Bioinformatics 31, 2382–2383 (2015).

82. Durand, N.C. et al. Juicer Provides a One-Click System for Analyzing Loop-Resolution Hi-C Experiments. Cell Syst. 3, 95–98 (2016).

83. Korsunsky, I., Nathan, A., Millard, N. & Raychaudhuri, S. Presto scales Wilcoxon and auROC analyses to millions of observations. Preprint at bioRxiv 10.1101/653253 (2019).

84. Fornes, O., et al. JASPAR 2020: update of the open-access database of transcription factor binding profiles. Nucleic Acids Res. 48, D87–D92 (2020).

85. Tan, G. & Lenhard, B. TFBSTools: an R/bioconductor package for transcription factor binding site analysis. Bioinformatics 32, 1555–1556 (2016).

86. Seitzer, P., Wilbanks, E.G., Larsen, D.J. & Facciotti, M.T. A Monte Carlo-based framework enhances the discovery and interpretation of regulatory sequence motifs. BMC Bioinformatics 13, 317 (2012).

