## Supplementary Figures for "Predicting enhancer-gene links from single-cell multi-omics data by integrating prior Hi-C information"

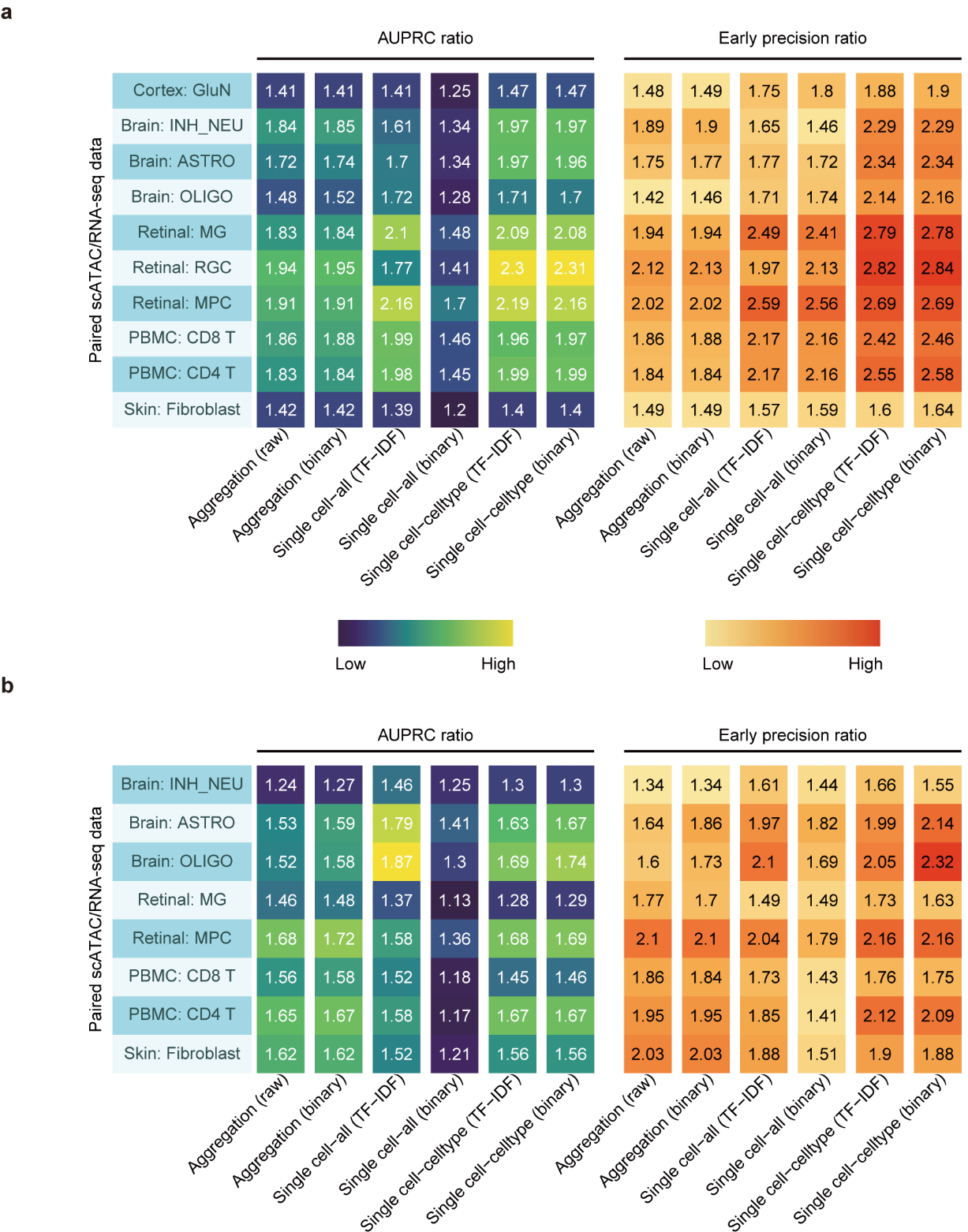

**Supplementary Fig. 1. Evaluation of data preprocessing methods for enhancer-gene link predictions in SCEG-HiC**

**a-b**, Performance across multiple human cell types was evaluated by AUPRC ratio (left) and EPR (right), and validated using cell type-specific Hi-C **(a)** and eQTL data **(b)**, respectively. Three main factors were considered for comparison: aggregation vs. single-cell retention, scATAC-seq binarization vs. non-binarization, across all cells vs. within a specific cell type. For single-cell retention approach, TF-IDF normalization was applied without scATAC-seq binarization.

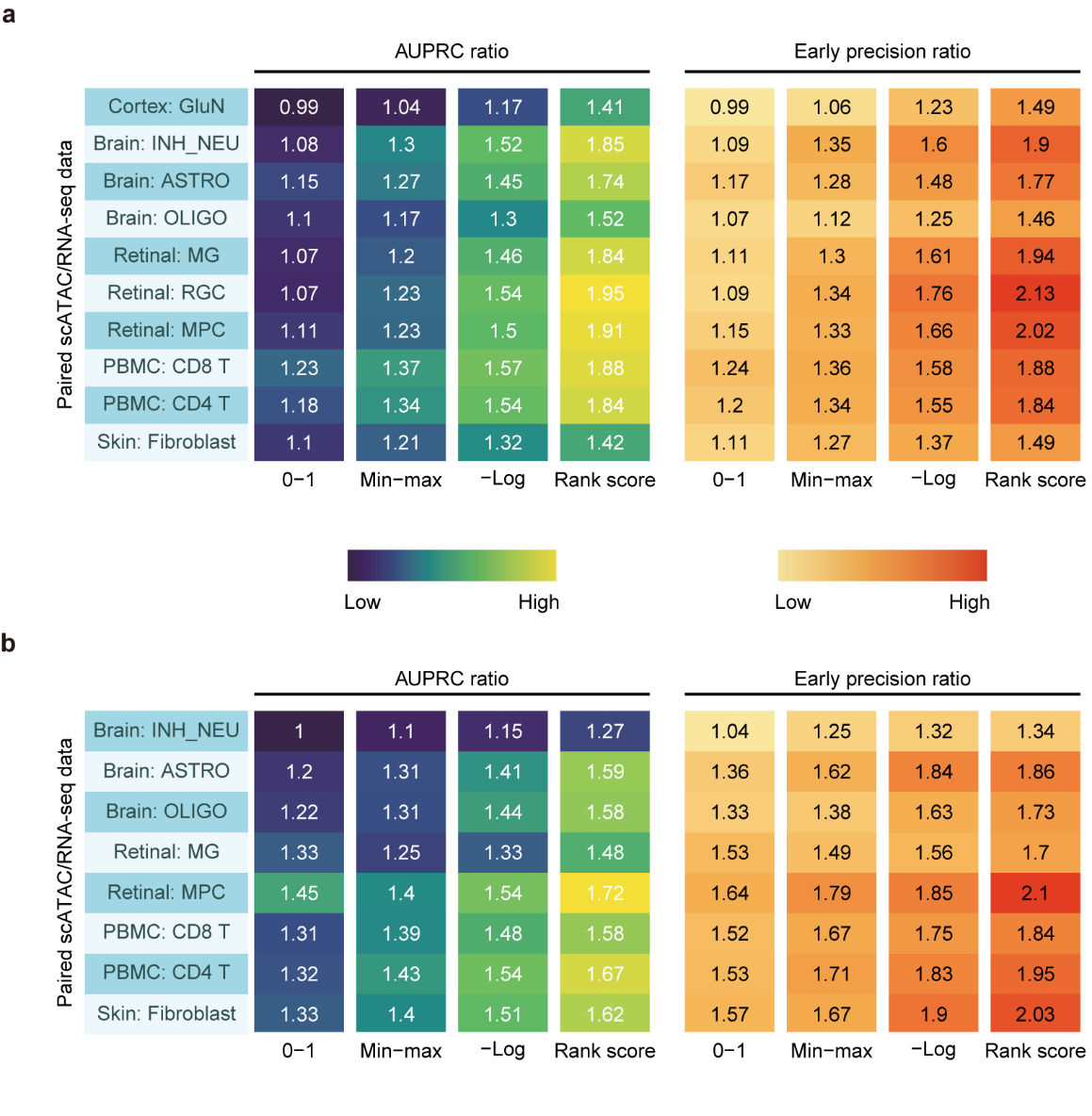

**Supplementary Fig. 2. Evaluation of bulk average Hi-C normalization methods for enhancer-gene link predictions in SCEG-HiC.**

**a-b**, Performance across multiple human cell types was evaluated by AUPRC ratio (left) and EPR (right), and validated using cell type-specific Hi-C **(a)** and eQTL data **(b)**, respectively. Four methods were considered for comparison, including 0-1 binarization, min-max normalization, -log transformation, and rank score.

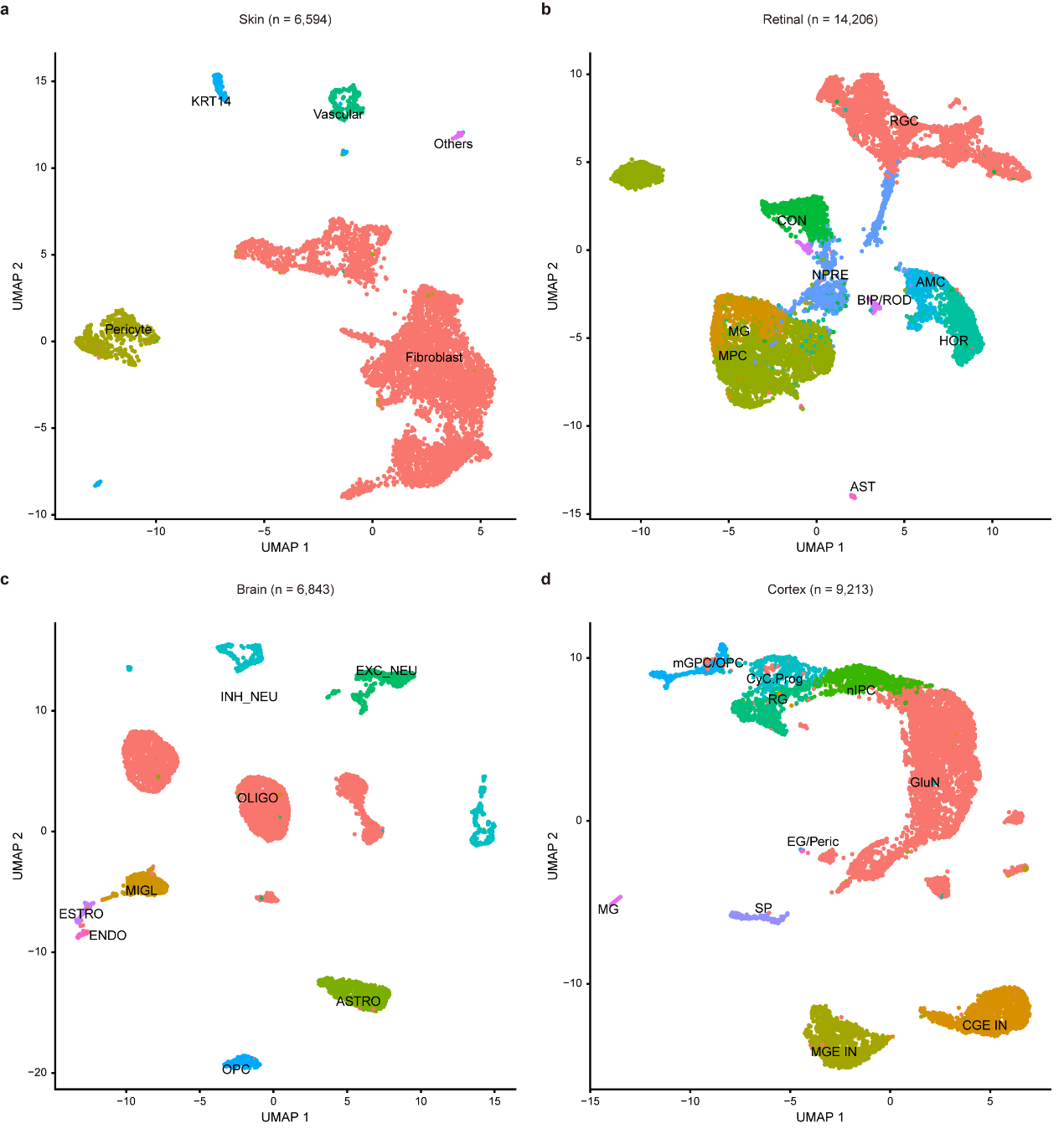

**Supplementary Fig. 3. Visualization of paired human single-cell multi-omics datasets.**

**a-d**, UMAP embeddings of single cells from skin stromal cells **(a)**, fetal retinal tissue **(b)**, brain gray matter **(c)**, and developing cerebral cortex **(d)** (*n* = cell number). Cell types were assigned based on known markers. AMC, amacrine cells; AST, astrocytes; BIP/ROD, bipolar and rod photoreceptor cells; CON, cone photoreceptors; HOR, horizontal cells; MPC, multipotent progenitor cells; NPRE, neurogenic precursors; MG, muller glia; RGC, retinal ganglion cells. ENDO, endothelial cells; ASTRO, astrocytes; EXC_NEU, excitatory neurons; INH_NEU, inhibitory neurons; MIGL, microglia; OLIGO, mature oligodendrocyte; OPC, oligodendrocyte precursor cell. RG, radial glia; CyC.Prog, cycling progenitors; mGPC/OPC, multipotent glial progenitor cell/oligodendrocyte progenitor cell; nIPC, neuronal intermediate progenitor cell; GluN, glutamatergic neuron; CGE IN, caudal ganglionic eminence interneuron; MGE IN, medial ganglionic eminence interneuron; EC, endothelial cell; MG/Peric, microglia/Pericytes; SP, subplate.

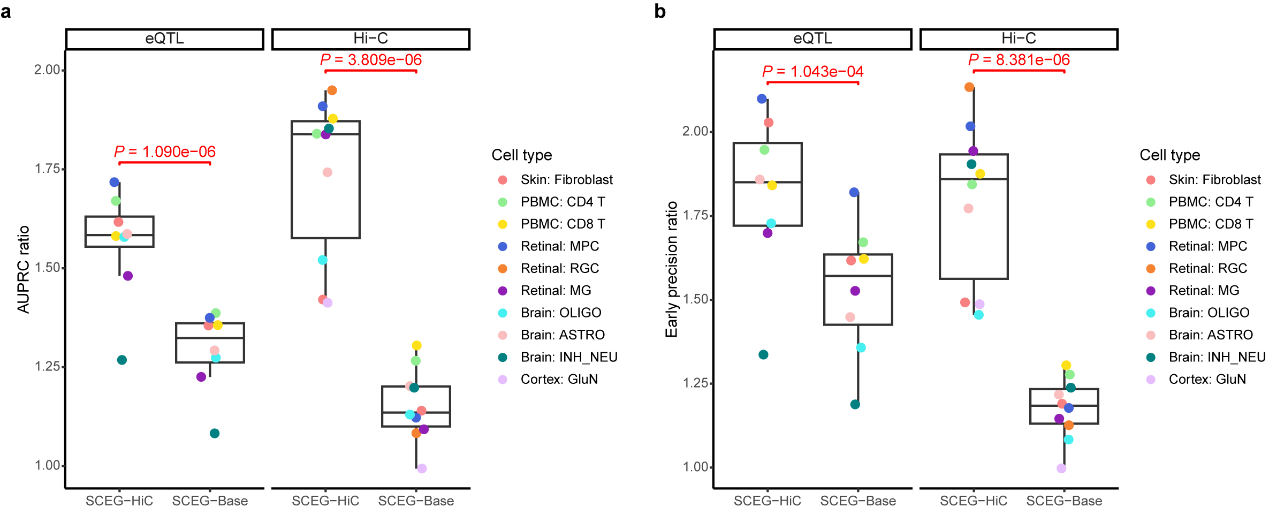

**Supplementary Fig. 4. Incorporating bulk average Hi-C data improves prediction accuracy of enhancer-gene links.**

**a-b**, Boxplots showing performance comparison between SCEG-HiC and SCEG-Base across various human cell types, measured by AUPRC ratio **(a)** and EPR **(b)**. SCEG-Base was constructed using the same model architecture as SCEG-HiC but with bulk average Hi-C data excluded. Validation was performed using cell type-specific Hi-C and eQTL data, respectively. Each dot represents a human cell type. *P*-values were calculated using two-sided paired *t*-tests.

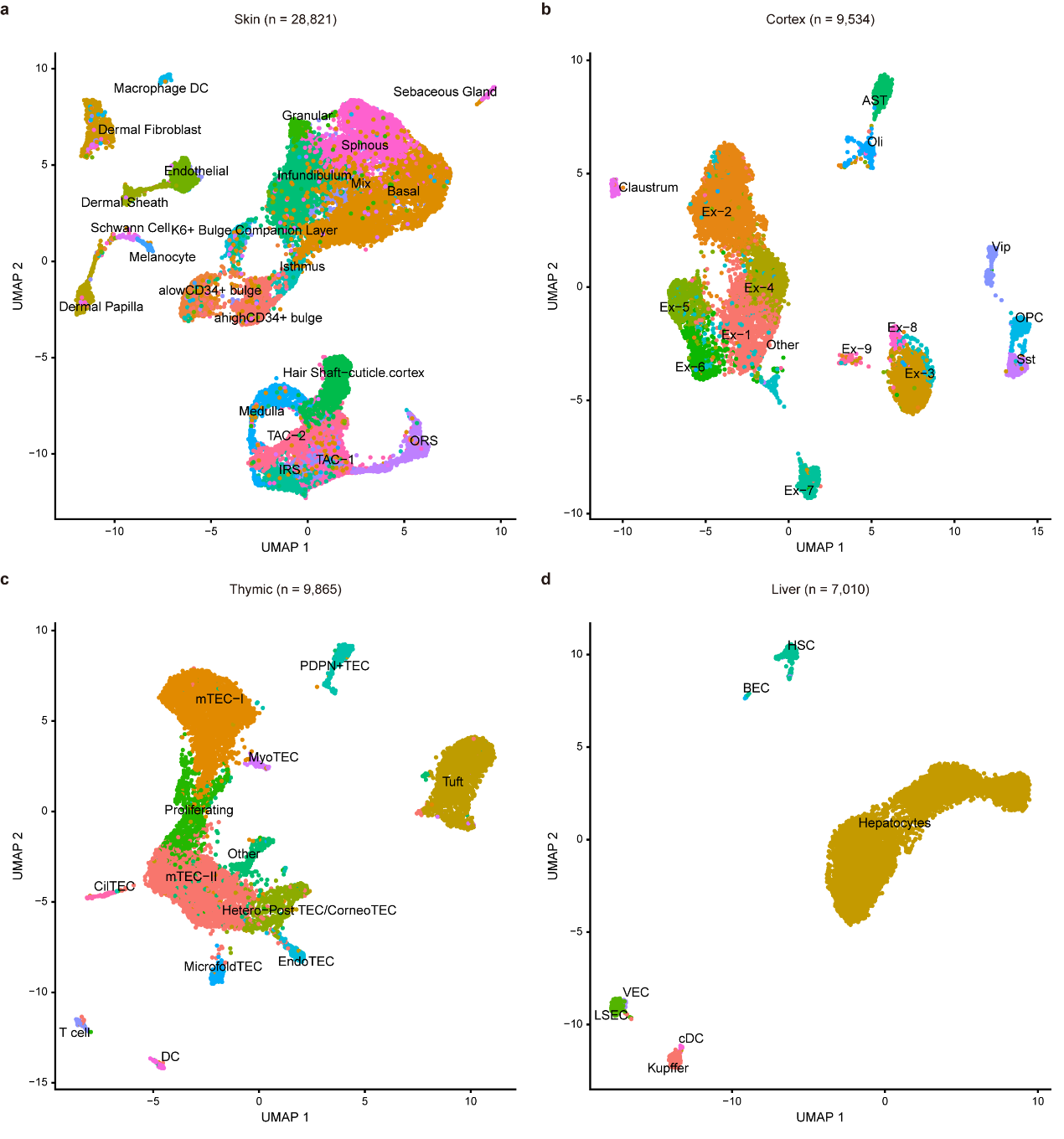

**Supplementary Fig. 5. Visualization of paired mouse single-cell multi-omics datasets.**

**a-d**, UMAP embeddings of single cells from mouse skin **(a)**, cerebral cortex **(b)**, thymic epithelial cells **(c)**, and liver **(d)** (*n* = cell number). Cell types were assigned based on known markers or annotations from the original publications. TACs, transit-amplifying cells; IRS, inner root sheath; ORS, outer root sheath. Ast, astrocytes; Ex, excitatory neurons (multiple subtypes); OPC, oligodendrocyte progenitor cells; Sst, Vip, inhibitory neuron subtypes; Oli, oligodendrocytes. TEC, thymic epithelial cell; mTEC-I, mTEC-II, well-characterized medullary TEC; Proliferating, transit-amplifying TECs; MyoTECs, myocyte-like TECs; CilTECs, ciliated-like TECs; CorneoTECs, corneocyte-like TECs; endoTECs, endocrine-like TECs; microfoldTECs, M-cells-like TECs; Hetero-post TEC, heterogeneous TECs; DC, dendritic cells. BEC, biliary epithelial cells; cDC, conventional dendritic cells; HSC, hepatic stellate cells; VEC, vascular endothelial cells; LSEC, liver sinusoidal endothelial cells.

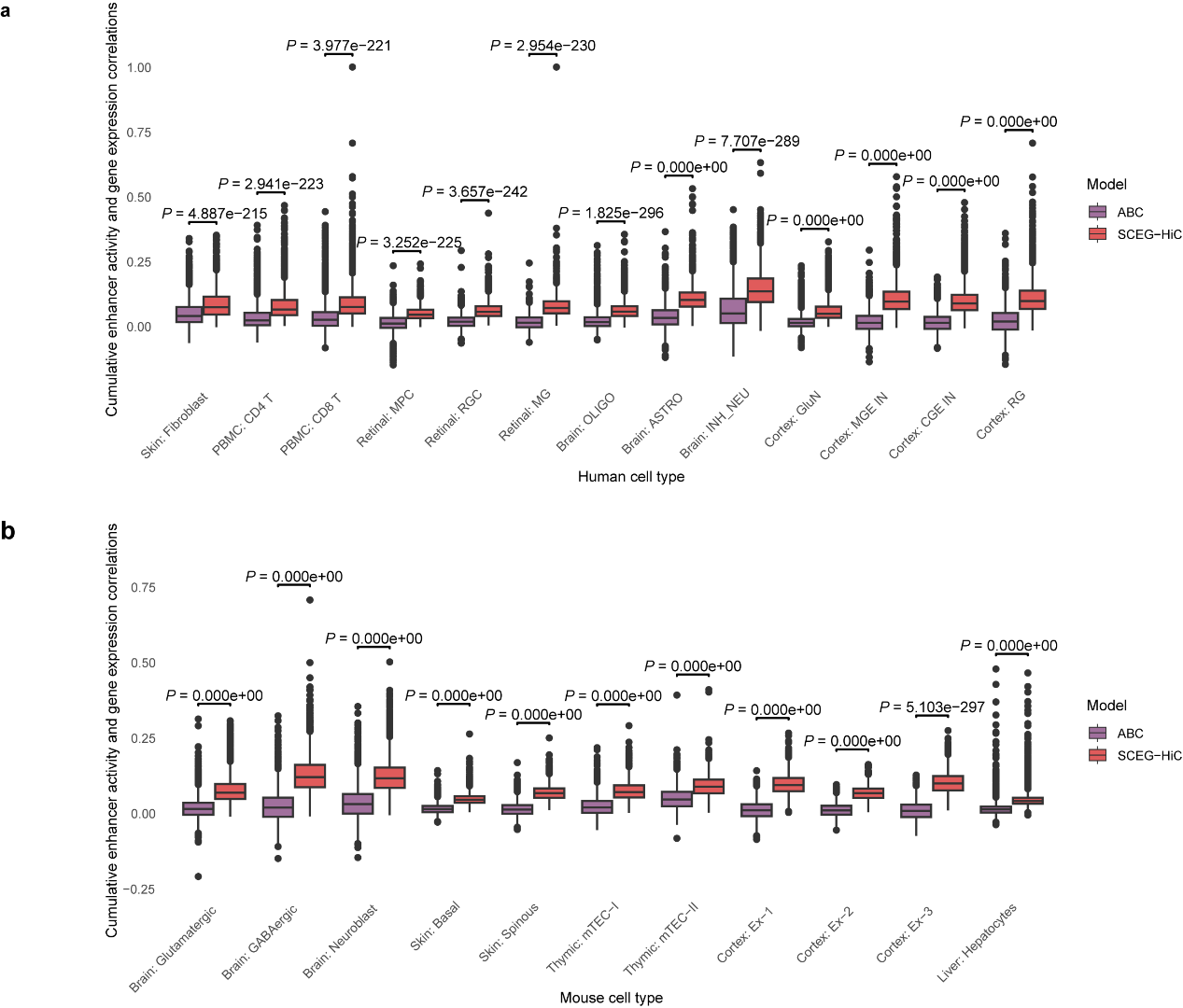

**Supplementary Fig. 6. Comparison of cumulative enhancer activity and gene expression correlations between SCEG-HiC and ABC model.**

**a-b**, Boxplots showing the Spearman correlation between predicted cumulative enhancer activity and gene expression for highly variable genes across human **(a)** and mouse **(b)** cell types. Each dot represents an individual gene. *P*-values were calculated using two-sided paired Wilcoxon tests.

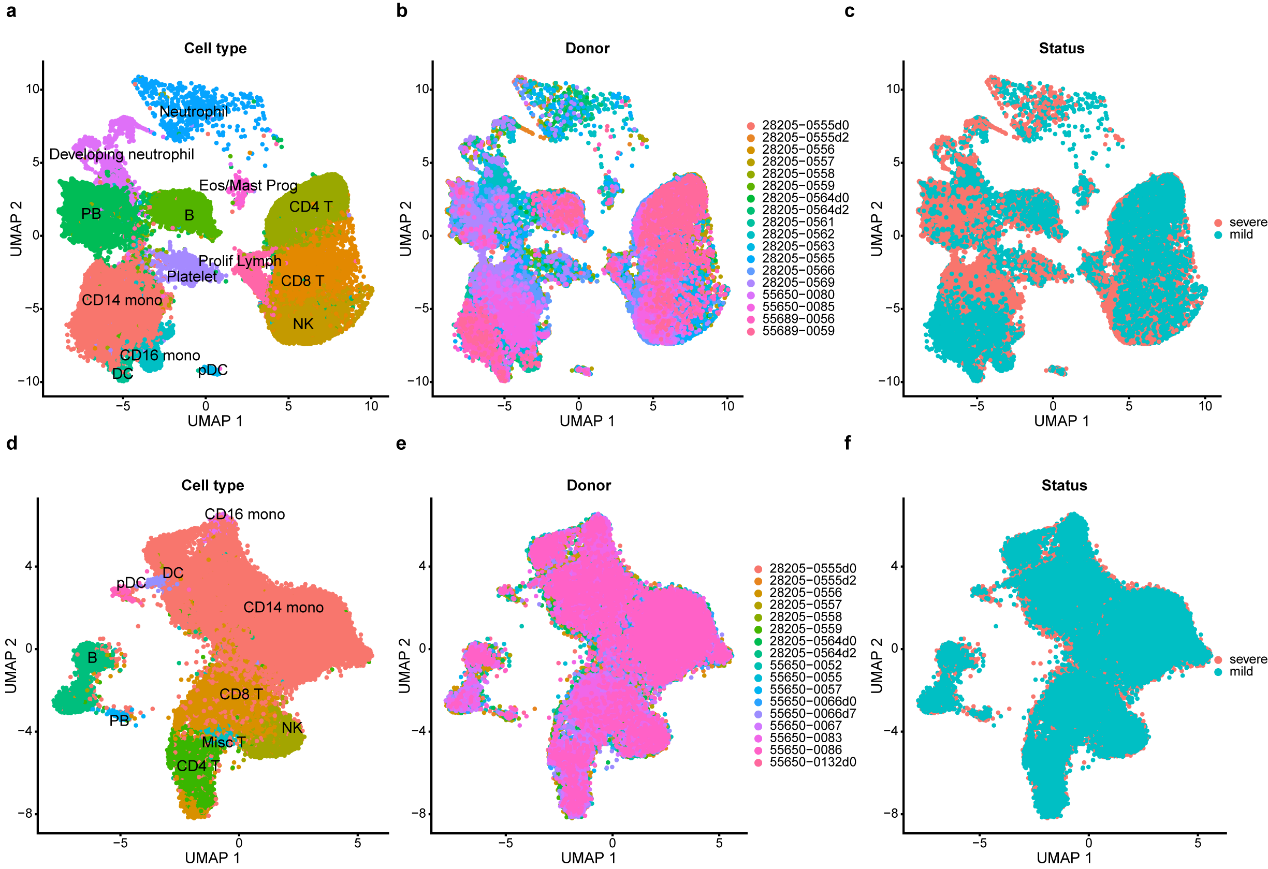

**Supplementary Fig. 7. Visualization of unpaired PBMC scRNA-seq and scATAC-seq datasets from COVID-19 patients.**

**a-c**, UMAP embeddings of the scRNA-seq dataset, colored by cell type **(a)**, donor **(b)**, or status **(c)**, respectively. Cell types were assigned based on annotations from the original publications. CD4 T, CD4+ T cell; CD8 T, CD8+ T cell; CD16 mono, CD16+ monocyte; CD14 mono, CD14+ monocyte; NK, natural killer cell; DC, dendritic cell; pDC, plasmacytoid dendritic cell; PB, plasmablast; Eos, eosinophils; Prog, progenitor; Prolif Lymph, proliferating lymphocytes.

**d-f**, UMAP embeddings of the scATAC-seq dataset, colored by cell type **(d)**, donor **(e)**, or status **(f)**, respectively. Cell types were assigned via label transfer from paired multi-omics reference data. Misc T, miscellaneous T cell, including mucosal-associated invariant T (MAIT), gamma-delta T (gdT), double-negative T (dnT), and regulatory T (Treg).

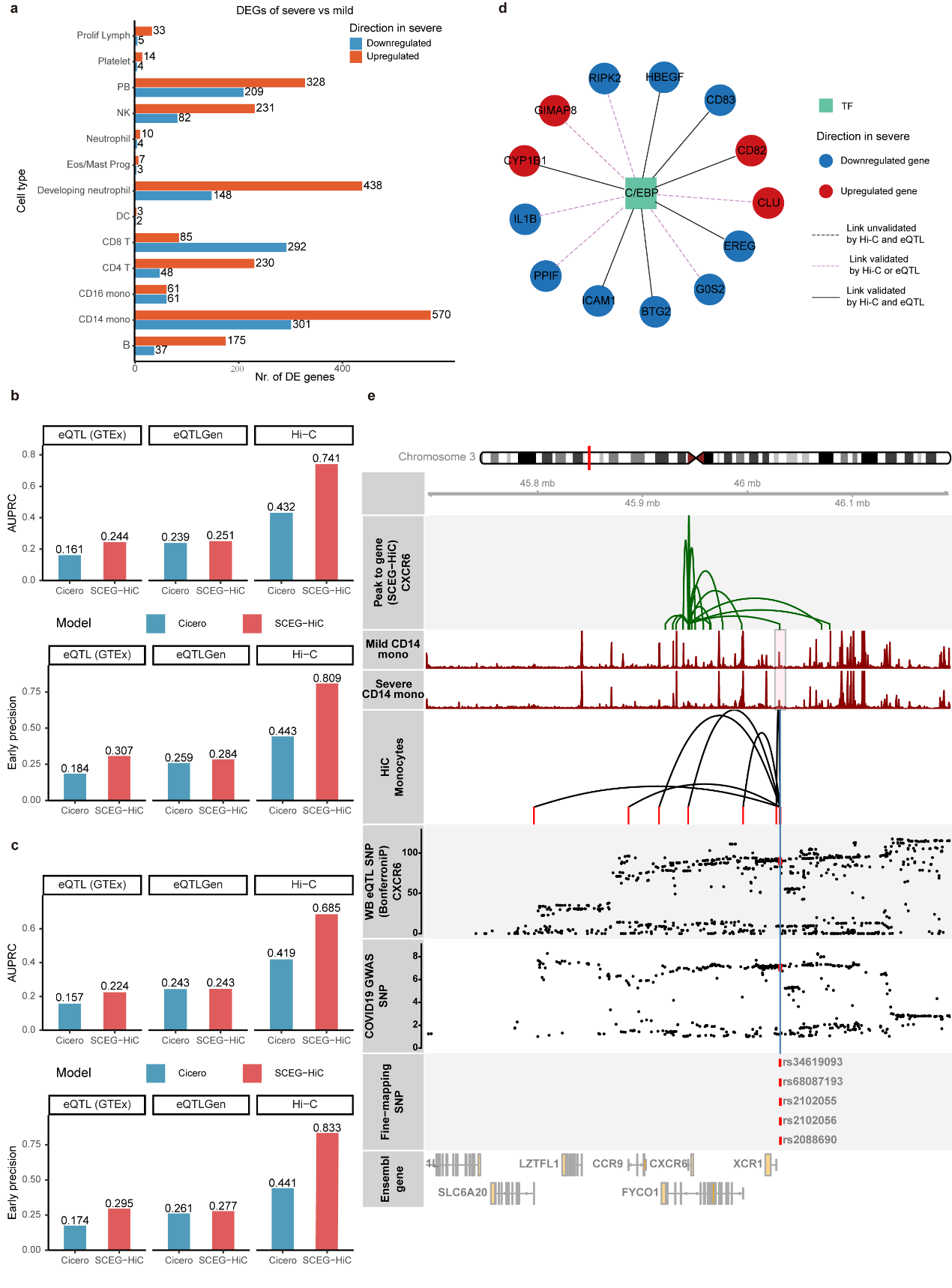

**Supplementary Fig. 8. Supplementary information for COVID-19 single-cell data analysis.**

**a**, Bar plots showing the number of significant DEGs identified in each cell type. Differential expression analysis was performed to compare severe vs mild individuals via Wilcoxon ranked-sum test.

**b-c**, Bar plots comparing performance between SCEG-HiC and Cicero in mild **(b)** and severe **(c)** CD14+ monocytes, respectively. The performance was measured by AUPRC (top) and EP (bottom), validated using CD14+ monocyte Hi-C, and whole blood eQTL data from GTEx and eQTLGen.

**d**, Subnetwork of the GRN between C/EBP TF families and target genes, inferred from TF motif enrichment analysis of SCEG-HiC-predicted enhancer-gene links.

**e**, SCEG-HiC prediction of fine-mapped SNPs linked to *CXCR6*. Integration of SCEG-HiC-predicted enhancer-gene links, scATAC-seq signals, CD14+ monocyte Hi-C interactions, whole blood eQTL SNPs, and COVID-19 GWAS SNPs around rs34619093, rs68087193, rs2102055, rs2102056 and rs2088690.
